# Molecular and cellular dynamics of measurable residual disease progression in myelodysplastic syndromes

**DOI:** 10.1101/2025.10.17.682671

**Authors:** Yu-Hsiang Chen, Timothy Bi, Minhang Xu, Jaehyun Lee, Wesley S. Lu, Mengchuan Zhuang, Alejandro Marinos Velarde, Dustin McCurry, Hui Yang, Gheath Al-Atrash, Gabriela Rondon, Elizabeth Shpall, Fatima Zahra Jelloul, Frances N. Cervoni-Curet, Sassine Ghanem, Celine Kerros, Priya Koppikar, Jeffrey Molldrem, Guillermo Garcia-Manero, Ankit Patel, Pavan Bachireddy

## Abstract

Cancer relapse after treatment invariably proceeds from the persistence and progression of measurable residual disease (MRD), an ultrasmall malignant population. Despite the poor prognostic impact established by increasingly sensitive MRD detection methods, the molecular pathways defining MRD remain unknown. To identify the unique features of MRD and the molecular forces shaping its progression, we performed single-cell multi-omic profiling on DNA, RNA and protein layers for a longitudinal patient cohort with relapsed myelodysplastic syndromes (MDS) after stem cell transplantation (SCT), the only curative modality for MDS. We provide a comprehensive molecular portrait of MRD cells with novel markers, shared across genetically heterogeneous patients. MDS relapse after SCT manifested universally with marked phenotypic evolution. Genotype and phenotype analyses revealed MRD progression as a dynamic, evolutionary process rather than a static expansionary one, driven by both subclonal sweeping and cell state transitions. Malignant cells adapted to infiltrating T cells by rewiring IFN-γ responses to activate a key immunoevasive pathway. Our study demonstrates the power of longitudinal, single-cell multi-omic analysis for identifying, tracking, and understanding MRD cells, opening new avenues to target MRD persistence and progression.

## Main Text

Myelodysplastic syndromes (MDS) are a family of stem cell disorders caused by somatic mutations that impair the differentiation of hematopoietic stem cells into mature progeny, leading to clonal expansion (*1–3*). Approximately 30% of MDS patients evolve into acute myeloid leukemia (AML), which carries a poor prognosis with a median survival of less than one year (*4–6*). Allogeneic hematopoietic stem cell transplantation (SCT) remains the only curative treatment for higher-risk MDS (HR-MDS) patients, who are at increased risk of progression to AML (*7–9*). However, relapse is still frequently observed after SCT despite the anti-leukemic effects of conditioning regimens and the graft-versus-leukemia (GvL) response (*6, 10*). PostSCT relapse is associated with dismal outcomes, with few effective salvage options and a median survival of 4.7 months (*6*), underscoring the need for predictive markers and post-transplant monitoring strategies.

In recent years, one major predictor of MDS relapse after SCT that has emerged is the presence of measurable residual disease (MRD) at complete remission—small numbers of cancer cells that persist after transplant (*11, 12*). Multiparameter flow cytometry, digital droplet PCR (ddPCR), and next-generation sequencing (NGS) have been used to detect MRD cells at detection limits ranging from 1 in 1,000 to 1 in a million (*13–16*). However, in addition to lacking technical harmonization, these techniques offer limited insights beyond quantitative MRD assessments and cannot characterize their molecular profile, leaving critical gaps in our understanding of how these cells persist and drive relapse. Unraveling their biological features will establish a mechanistic foundation for revoking MRD progression and provide broadly applicable principles for MRD biology across myeloid malignancies.

In this study, we developed a novel single-cell multi-omic workflow, termed CARAMEL-seq, which integrates scRNA-seq, scATAC-seq, surface protein profiling, and lineage-informing mutations to 1) capture rare malignant MRD cells, 2) characterize their unique molecular features, and 3) track phenotypic and clonal changes of MDS cells from SCT through complete remission to relapse using longitudinal sampling. This framework provides a powerful approach to uncover mechanisms of residual cancer cell persistence and evolution to drive relapse.

### Donor–recipient dynamics across cell compartments after SCT

*In vitro* and *in vivo* MDS models offer limited ability to capture the genomic and phenotypic heterogeneity and behavior of MDS cells, especially under treatment pressures (*17, 18*). To address these constraints in the context of MRD and to investigate dynamic changes in MDS cell populations after SCT, we utilized bone marrow mononuclear cell (BMMC) samples collected and cryopreserved before SCT (preSCT), during complete remission (CR), and at relapse (Fig. 1A-B). BMMCs were sorted by flow cytometry into CD34 stem/progenitor and CD34 immune cell fractions for compartment-specific analyses. Both fractions were analyzed using CARAMEL-seq to capture **C**hromatin **A**ccessibility (scATAC-seq), transcriptomic (sc**R**N**A**-seq), **M**itochondrial variant–based clonal mutations for lineage and clonal inference, and **E**pitope **L**abeling (surface-protein expression) information (Fig. S1). Our study cohort included 13 patients, comprising 10 individuals who relapsed post-transplant (postSCT) and 3 who remained relapse-free (Table S1 and S2). None of the patients were treated with maintenance therapy after SCT, presenting a unique opportunity to study the natural history of MRD progression (*19*). From 52 samples, 192,149 high-quality CD34 and 138,786 CD34 cells were attained for downstream analysis after filtering (Fig. S2A-B).

**Fig. 1.**
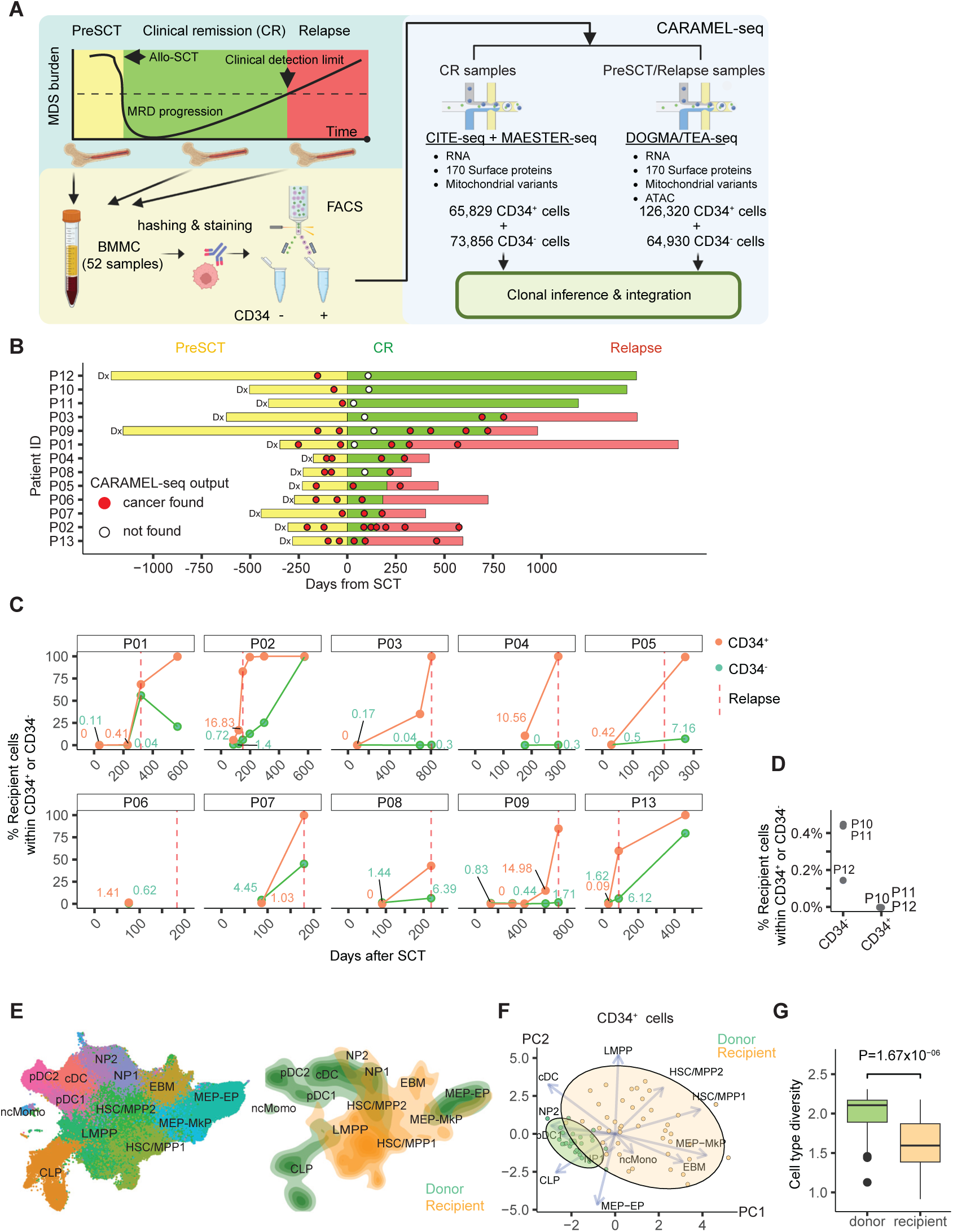
**Single-cell multi-modal profiling of CD34**D **and CD34**D **bone marrow cells during SCT and relapse.** (A) Schematic illustration of study design. Bone marrow mononuclear cells (BMMC) from pre-transplant (preSCT; yellow), clinical remission (CR; green), and relapse (red) timepoints were FACS-sorted into CD34 and CD34 fractions. Single-cell multi-omics profiling was performed using CITE-seq + MAESTER-seq for samples from CR; and DOGMA/TEA-seq for samples from preSCT and relapse to capture RNA, 170 surface proteins, mitochondrial variants, and ATAC data. (B) Timeline of sampling relative to SCT for each patient. Bars indicate disease status classified as preSCT (yellow), CR (green), or relapse (red), beginning from diagnosis (Dx). Circles denote presence (red), or absence (white) of malignant cells detected by single-cell analysis. (C) Longitudinal quantification of recipient-derived cell fractions in CD34 and CD34 compartments among relapsing patients after SCT, with relapse timepoints indicated by vertical, dotted red lines. Labels indicate the percentage of recipient-derived cells. (D) Recipient-derived cell percentages in CD34 and CD34 compartments at remission in patients without relapse. (E) UMAP projections of donor-derived and recipient-derived cells colored by hematopoietic cell type annotation (left) or donor/recipient density (right). (F) PCA of donor and recipient cells highlighting shifts in lineage bias in recipient cells with shaded ellipses indicate 95% confidence intervals. Each circle represents a sample. (G) Boxplot comparing cell type diversity (Shannon index) between donor and recipient-derived cells in each sample. Statistical significance was calculated by two-sided Wilcoxon test.

Following SCT, donor- and recipient-derived cells may coexist within the bone marrow, a phenomenon known as chimerism. To deconvolve donor-recipient origin at the single-cell level, we employed Souporcell genotype demultiplexing (*20*) to assign cells as donor- or recipient-derived, achieving an estimated assignment error rate of 0.06% (Fig. S2C). In nine of ten patients who relapsed, recipient-derived cells were detectable at CR timepoints in both the CD34 stem/progenitor compartment (0.09%–16.83%) and the CD34 immune compartment (0.04%– 4.45%) (Fig. 1C). In contrast, in the three patients who remained in CR, recipient-derived cells could only be found at orders of magnitude lower levels within the CD34 fraction (0.14%– 0.45%), and they were undetectable in the CD34 fraction (Fig. 1D). Between CR and relapse, recipient-derived CD34 cells emerged earlier and more rapidly than CD34 cells across all patients, suggesting that re-growth kinetics differ markedly between progenitors and progeny, and that the resurgence of MDS stem/progenitor cells drives morphological relapse. These findings underscore the value of enriching and monitoring the CD34 compartment, which typically constitutes less than 2% of bone marrow cells, as an early molecular indicator of relapse risk following transplantation (*16*).

To identify variations in differentiation and lineage allocation between donor- and recipient-derived hematopoiesis, we next examined the cell-type composition within these CD34 populations. We first constructed a reference multimodal map of normal CD34 hematopoietic stem and progenitor cells (HSPCs) using donor-derived cells, with cluster annotations based on RNA and surface-protein profiles (Fig. S3). This analysis revealed 13 populations, including two hematopoietic stem and multipotent progenitor (HSC/MPP) clusters, with HSC/MPP1 showing a more long-term stem-like state characterized by lower pseudotime values and higher expression of CD90 and *HLF* relative to HSC/MPP2, which represented a more differentiated state (Fig. 1E and S3B-D).

Recipient cells were mapped back to this reference to identify their corresponding normal counterparts, achieving a median mapping score of 78% compared with 98% for donor cells (Fig. S4), underscoring the molecular divergence of recipient cells from normal HSPCs. Overall, recipient-derived cells were enriched in HSC/MPPs and showed reduced representation of common lymphoid progenitors (CLP) compared to donor-derived cells, indicating a recipient bias toward primitive progenitors (Fig. 1E and S5). Indeed, the clinical multiparametric flow cytometry (MFC) observations indicated that >80% of our patients exhibited CD38 decrease or negativity in the cohort (Table S1), consistent with the CD38 ^/dim^ phenotype of HSC/MPPs. Analysis of cell-type composition across samples revealed a clear separation between donor- and recipient-derived populations based on principal component analysis (PCA), with greater inter-sample variation observed among recipient-derived populations (Fig. 1F). However, recipient-derived cells exhibited significantly lower cell-type diversity within individual samples compared to donor-derived cells (*p*=1.67×10) (Fig. 1G). These results indicate that, *within* each patient, recipient cells display lower cell-type diversity compared to donor cells, while *across* patients, recipient cells display greater compositional diversity than donor cells. In summary, we characterized the differentiation dynamics and differences between donor- and recipient-derived cells. Furthermore, we recovered recipient-derived cells at low levels during CR, which likely represent residual malignant cells that ultimately drive morphologic relapse.

### Identification of ultra-low-level MRD at flow cytometry–negative CR timepoints

Although residual recipient cells after SCT are generally considered malignant due to their resistance to conditioning regimens and survival advantage over normal hematopoietic cells, the persistence of non-malignant (“normal”) recipient cells is not rare (*16, 21, 22*). We therefore sought to distinguish malignant from normal recipient populations and to quantify their relative proportions during CR. 17 out of 19 CR timepoints showed no evidence of MDS by clinical, standard-of-care flow cytometry, and all exhibited 100% donor chimerism in the myeloid cells (Table S2; Methods). Patients with MDS exhibit diverse cytogenetic, genetic, and immunophenotypic alterations, resulting in a lack of universal markers which can reliably distinguish MRD from normal recipient cells. Nevertheless, the multi-modal measurements provided by CARAMEL-seq, in conjunction with the aberrant immunophenotypes known for each patient, allowed us to comprehensively interrogate each modality to resolve malignant versus normal cells (Fig. 2A). In patients P01 and P09, we identified clonal, myeloid-specific mitochondrial variants found in our ATAC data which delineated malignant recipient-derived cells with high specificity (Fig. 2B-C and fig. S6A-C). Additionally, approximately 50% of MDS patients harbor somatic copy number variations (CNVs), while 5–10% of male patients exhibit loss of chromosome Y (chrY) (*23*). Our cohort included 7 of 13 patients with clinically documented CNVs and 2 patients with loss of chrY. We were able to recapitulate these genomic events by inferring CNV profiles from the RNA modality (Fig. 2D, S6D, and S7), confirming concordance between single-cell CNV inference and clinical cytogenetic data and identifying MRD cells in these patients. Finally, aberrant expression of markers typically restricted to mature hematopoietic cells (e.g. CD5, CD7, CD9, and CD56) on HSPCs is common and widely seen in AML/MDS. In our cohort, only P02 and P13 were documented to have an aberrant immunophenotype with overexpression of CD5 and CD56; our single-cell data confirmed that CD5 overexpression was limited to these individuals and discriminated MRD from non-malignant recipient cells (Fig. 2E). Lastly, in patient P13, single-cell UMAP and low-resolution clustering clearly separated most donor and recipient cells, with only a small fraction of recipient cells appearing in the donor-enriched cluster. This donor-dominant cluster was used as a reference for normal cells to distinguish malignant and normal recipient-derived populations (Fig. S8). These results highlight the utility and necessity of leveraging multimodal features from CARAMEL-seq to identify malignant cells across MDS patients with diverse genetic and phenotypic backgrounds.

**Fig. 2.**
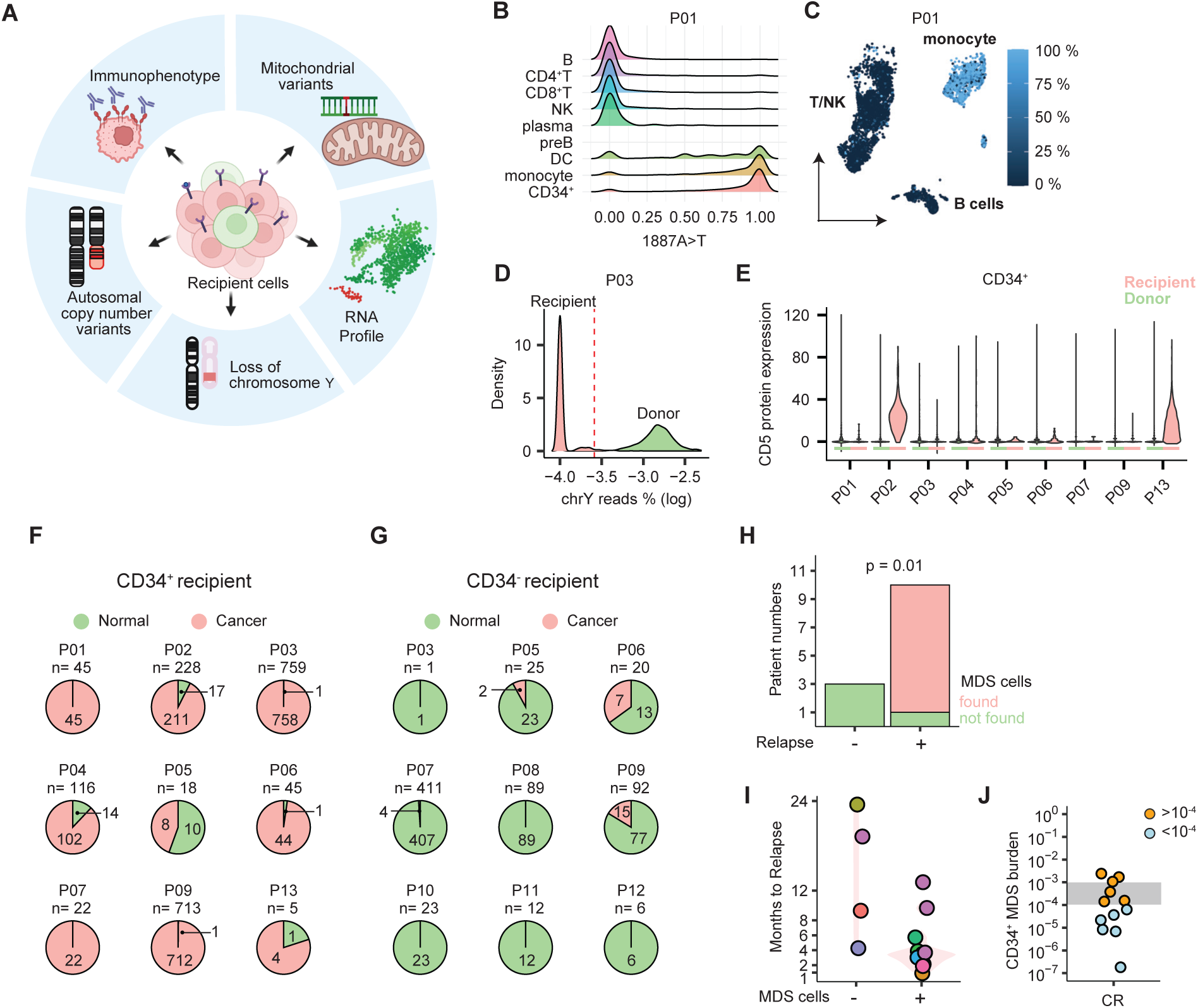
Multi-modal identification of residual malignant recipient-derived cells at complete remission. (A) Schematic illustration of CARAMEL-seq features used to distinguish malignant from normal recipient-derived cells at CR. (B) Density distributions of the 1887A>T mitochondrial variant allele frequency in patient P01 across different cell type compartments. (C) Allele frequency of the 1887A>T in patient P01 projected onto CD34 UMAP embeddings of single-cell transcriptomes. (D) Density plot of log-transformed chromosome Y read percentages in individual cells in patient P05, with the red dashed line indicating the threshold used to identify loss of chromosome Y events. (E) CD5 protein expression across patients, colored by donor or recipient source at CR. (F–G) Pie charts showing classification of recipient-derived cells in CD34 (F) and CD34 (G) compartments at CR, with proportions of malignant (pink) and normal (green) cells across patients. (H) Numbers of patients with or without relapse stratified by detection of malignant CD34 recipient-derived cells at CR. (I) Timing of CR sampling relative to relapse, stratified by detection of malignant CD34 recipient-derived cells at CR. Different colors represent individual patients. (J) Distribution of CD34 cancer burden at CR among relapsing patients. Gray shading indicates the sensitivity threshold of flow cytometry; blue points represent values below 10^-4^, and orange points represent values > 10^-4^.

CARAMEL-seq enabled us to successfully distinguish recipient-derived normal from malignant cells from MRD-negative (by MFC) CR timepoints (Fig. 1B); therefore, we use the term “ultra-low-level MRD (uMRD)” to denote their detection. We then sought to determine the proportion of malignant and non-malignant cells constituting recipient-derived CD34 and CD34 compartments at these uMRD timepoints. Within CD34 fractions, recipient-derived cells from CR samples in relapsing patients were predominantly classified as malignant, ranging from 44% to 100% of the CD34^+^ compartment (Fig. 2F). In contrast, recipient-derived cells in the CD34 compartment were largely classified as normal, with malignant proportions ranging from 0% to 35% (Fig. 2G). These data suggest that the faster re-growth kinetics observed in the recipient CD34^+^ cells (Fig. 1D) are likely driven by the higher proportion of malignant cells overtaking that compartment. Moreover, they suggest that non-malignant CD34^+^ progenitors can survive and persist up to 4 months after high-intensity conditioning regimens.

Overall, no malignant cells were detected in either compartment among the three patients who did not relapse, whereas uMRD cells were detected in 9 of 10 patients who ultimately relapsed (*p*=0.01) (Fig. 2H), underscoring the predictive value of malignant cell detection within the CD34 compartment. uMRD was detectable from 1 to 13 months before morphologic relapse (Fig. 2I), indicating that CARAMEL-seq can identify malignant cells >1 year before relapse is detectable by flow cytometry. By integrating clinical estimates of total bone marrow CD34 prevalence with our measurements of CD34 malignant cell proportions, we measured CD34 uMRD frequencies ranging as low as 10^-7^, exceeding the typical sensitivity of current flow cytometry assays (<10). We also detected uMRD at frequencies as high as 10^-3^, indicating the additional discriminatory ability of our method to identify malignant cells even at frequencies well within the detectable range of conventional assays (Fig. 2J). Thus, by leveraging multiple modes of molecular measurements through CARAMEL-seq, we could discern (non)malignant, recipient-derived cells at uMRD, chart their relative dynamics through stem/progenitor (CD34) and progeny (CD34) compartments, measure their frequencies, and demonstrate their association with relapse.

### Unique molecular features of uMRD cells

Beyond merely detecting uMRD, we aimed to characterize these cells to deepen our understanding of MRD biology and gain insights into the mechanisms underlying disease persistence and progression. We first analyzed uMRD cell type composition at CR and unexpectedly found that uMRD cells were not a homogeneous group but rather comprised a set of varied phenotypes, with malignant cells mapping to a continuum of progenitor types (Fig. 3A). In addition, this progenitor pool composition differed markedly from donor-derived CD34 cells, with a notable enrichment for HSC/MPP-like states in uMRD populations (*p*=3.4×10^-5^), consistent across the genetically heterogeneous patient cohort (Fig. 3A-B). Given this enrichment and the fact that MDS is rooted in disrupted stem cell function (*24, 25*), we next aimed to identify protein and RNA features that discriminate malignant HSC/MPP-like cells from donor-derived HSC/MPPs at CR.

**Fig. 3.**
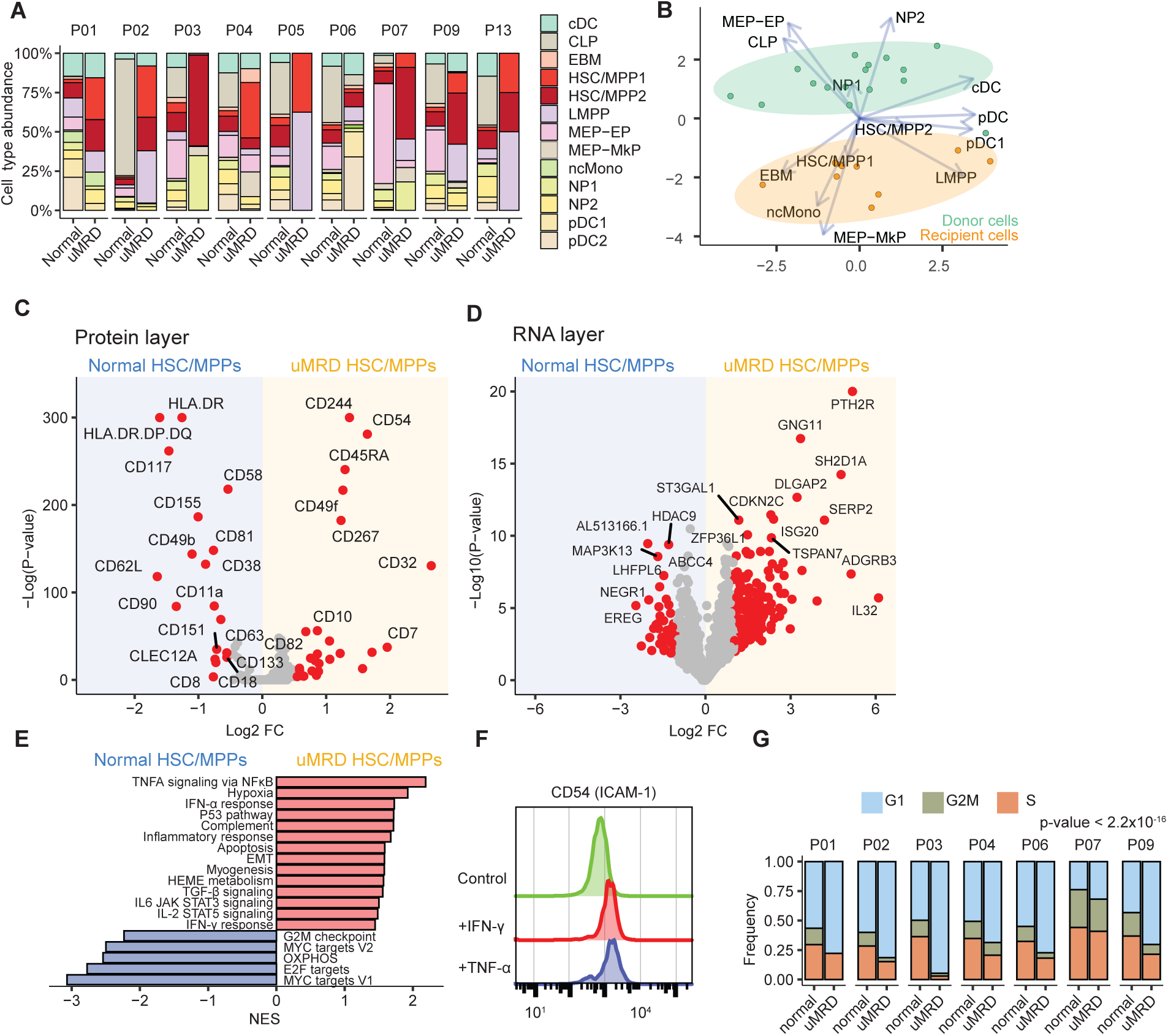
Distinct differentiation states, surface-protein profiles, and transcriptional programs in uMRD versus normal HSPCs. (A) Cell-type composition within CD34 populations for each patient between uMRD cells and normal HSPCs from donor cells. (B) PCA of cell-type composition in normal and recipient malignant compartments, with ellipses indicating 95% confidence intervals and arrows indicating the direction and strength of cell-type contributions (loadings) to the principal components. (C, D) Volcano plots of differential surface-protein (C) or gene (D) expression between uMRD and normal HSPCs, with selected proteins and genes labeled. (E) GSEA comparing RNA profiles of uMRD versus normal HSPCs. (F) Expression of CD54 (ICAM-1) on MDS-L cells under control conditions and after stimulation with IFN-γ and TNF-α. (G) Cell-cycle phase distributions for uMRD and normal HSPCs across patients; Mantel–Haenszel test p-value indicates that the G1/G0 phase is overrepresented in uMRD.

Analysis of 170 surface proteins in HSC/MPP-like uMRD cells revealed that, despite inter-patient heterogeneity, certain common alterations were observed. The most significantly increased surface markers in uMRD included CD54 (ICAM-1), observed in 8 of 9 patients, and CD244 (SLAMF4), elevated in 6 patients (Fig. 3C and Fig. S9A). Increased expression of CD54, a cell surface glycoprotein involved in inflammation, has been reported in MDS blasts with potential prognostic value (*26, 27*), though its functional role remains undefined. Similarly, the relevance of CD244 in MDS blasts has not been reported. We also detected overexpression of CD49f and GPR56, stemness-associated markers that have been reported in AML but have not previously been identified on MDS stem cells (*28–30*), as well as increased expression of CD45RA, a proposed leukemic stem cell (LSC) marker previously described in MDS (*31*). Among downregulated proteins in uMRD cells, reduced expression of HLA class II molecules— critical for T cell–mediated immune surveillance against abnormal HSPCs—was observed in 6 patients, along with downregulation of CD90 in 5 patients. These findings not only confirm known MDS markers but also reveal potential novel markers for improved identification and characterization of HSC/MPP-like malignant cells at uMRD.

We then compared the transcriptomic profiles of HSC/MPP-like uMRD cells to donor-derived HSC/MPPs. uMRD cells demonstrated significant upregulation of genes previously not associated with MDS including *GNG11*, *DLGAP2*, *ISG20*, and *LIMS1,* alongside previously reported genes such as *PTH2R* and *TGFB1*. *PTH2R* was recently identified as a potential marker of leukemic stem cells (LSCs), promoting survival through autophagy (*32*), and *TGFB1* is a key signaling node and therapeutic target in MDS (Fig. 3D and Fig. S9B)(*33–35*). Conversely, notable downregulated genes included *DPPA4*, *MAP3K13*, *CPED1*, *HDAC9*, and *MYC*. The downregulation of *MYC* contrasts with prior reports of *MYC* overexpression in MDS blasts during progression (*36*). This discrepancy may reflect a context-dependent role of *MYC*: in residual HSC/MPP-like populations, reduced MYC activity could (i) serve as an indicator of inflammatory stress (*37, 38*), and/or (ii) represent a distinct function of MYC in maintaining stemness or quiescence within MDS progenitors (*39*).

To examine pathway-level differences between malignant and donor-derived HSC/MPP cells, gene set enrichment analysis (GSEA) was performed (Fig. 3E). HSC/MPP-like uMRD cells showed positive enrichment of inflammatory and stress-related pathways, including TNF-α signaling via NF-κB, hypoxia, IFN-γ and IFN-α/β response, p53 pathway, and inflammatory response. We also utilized CytoSig (*40*) to infer cytokine signaling activities, as GSEA covers only a limited subset of cytokine signatures and does not resolve patient-specific activity. Notably, the increases in TGF-β, TNF-α, and IFN-γ signaling observed at CR were corroborated by CytoSig, which consistently revealed elevated activity of these cytokine pathways across patients (Fig. S10). The observed increase in inflammatory signaling across patients may explain the upregulation of CD54 surface protein, as CD54 can be induced by IL-1, TNF-α, and IFN-γ and plays a crucial role in immune responses and inflammation (*41–43*). We confirmed this inflammatory regulation in MDS *in vitro* by inducing CD54 expression in MDS-L cells via IFN- γ and TNF-α stimulation (Fig. 3F). Downregulated pathways in HSC/MPP-like uMRD cells included cell cycle and proliferation-associated pathways such as G2M checkpoint, MYC targets, E2F targets, and oxidative phosphorylation. Separately, cell cycle phase inference analysis further demonstrated a reduced fraction of cycling cells within the malignant population across every patient, consistent with the MYC downregulation and GSEA results we observed (Fig. 3G). Collectively, these results suggest that uMRD cells at CR have transcriptional signatures characterized by heightened inflammatory signaling and stress responses, alongside suppressed proliferative and metabolic activity.

### From SCT to relapse: cell state transitions of uMRD

We next sought to determine the relationship between uMRD and preceding preSCT or subsequent postSCT relapses: is uMRD a purely quantitative measure of residual/relapsed MDS, a distinct cell state or both? Similarly, is MRD progression a purely static expansionary process or a dynamic, evolutionary one? To compare cell states across preSCT, CR, and relapse timepoints, we individually labeled preSCT and relapse samples with hashtag oligos (HTOs) and processed them within the same experimental batch. These hashed samples, along with CR samples, were analyzed using MultiVI (*44*) for mosaic integration of RNA, protein and ATAC modalities. Batch correction was performed using the experimental batch as a covariate to minimize technical variation while preserving true biological differences, which can be obscured when using sample identity as a covariate. Among the six patients with multiple preSCT samples, we found that samples collected at different preSCT timepoints were phenotypically consistent with largely unchanged molecular states across various timepoints and therapies pre-SCT. In contrast, we observed significant cell state shifts of malignant HSC/MPPs from preSCT to relapse across all patients in the tri-modal-derived UMAP (Fig. 4A and S11), supported by a median Bray–Curtis similarity score of 0.2 (Fig. S12A-B). These data indicate that the observed cell state shifts are attributable to SCT itself rather than sampling variability and highlights SCT-induced phenotypic remodeling. Finally, we sought to identify the cellular source of this phenotypic change by measuring the dissimilarity between pre-SCT and relapse in the MultiVI latent space across cell types. Strikingly, we observed that the *degree* of cell state shift was the highest in the HSC/MPP compartment and decreased along differentiation trajectories in the myeloid and erythroid lineages across patients (Fig. 4B-D). These data suggest that MDS relapse after SCT involves extensive phenotypic evolution, characterized largely by molecular changes in the HSC/MPP compartment.

**Fig. 4.**
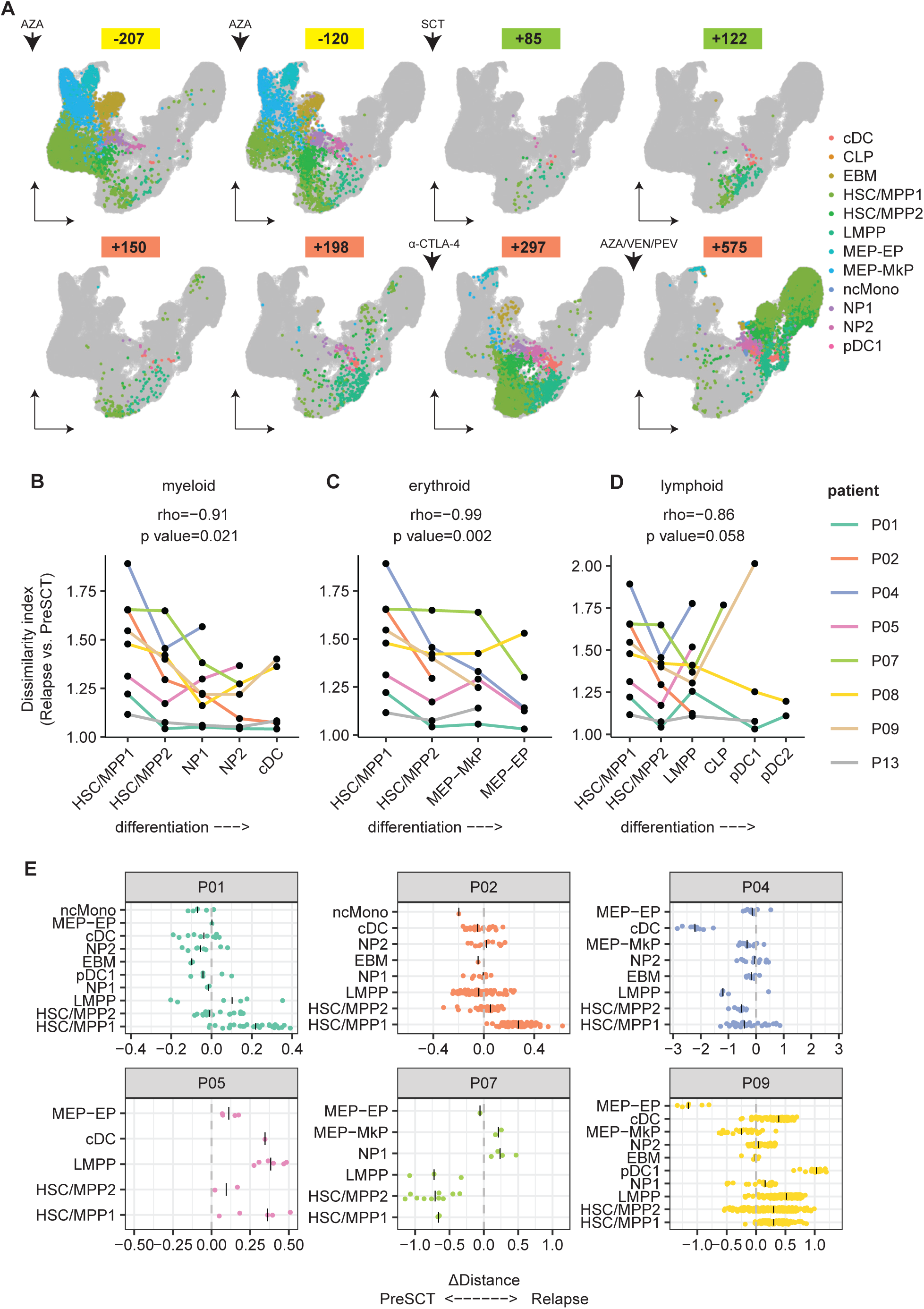
**Phenotypic transitions of uMRD CD34**D **cells from preSCT to relapse.** (A) UMAP projection from the trimodal-derived MultiVI latent space of uMRD CD34 cells in P02, colored by reference-mapped cell type annotations and stratified by sampling time. (B–D) Phenotypic dissimilarity scores between relapse and preSCT samples for each cell type stratified by myeloid (B), erythroid (C), and lymphoid (D) lineages across patients. Spearman correlation coefficients and p-values assessing the relationship between differentiation stage and deviation score across lineages. (E) Deviation scores of CD34 uMRD cells across annotated hematopoietic stem and progenitor types, comparing the phenotypic similarity of individual uMRD cells to preSCT versus relapse states. Positive values indicate greater similarity to relapse, while negative values indicate greater similarity to preSCT. AZA, Azacitidine; α-CTLA-4, ipilimumab; VEN, Venetoclax; PEN, Pevonedistat.

We next sought to determine whether uMRD cells more closely resemble the preSCT or relapse state. We calculated the cell distance between uMRD cells and both preSCT and relapse samples in the multiVI latent space, then determined deviation by subtracting these distances to assess relative similarity in the 6 patients with sufficient uMRD cells detected (i.e. n>10). In patients P01, P02, P05, and P09, uMRD HSC/MPP populations exhibited greater similarity to the relapse state than preSCT state (Fig. 4E). In contrast, in patient P04, who developed secondary AML (sAML) at relapse, most cell types were closer to the preSCT state. HSC/MPPs and LMPP-like cells in P07 likewise showed a preSCT bias. These results demonstrate both patient-level and cell-level heterogeneity in uMRD cells. For patients whose HSC/MPP cells showed a relapse bias at CR, the uMRD cells appeared already primed for relapse, adopting a relapse-like cell state at the time of CR sampling. Conversely, a preSCT bias in uMRD cells suggests that CR sampling occurred before phenotypic evolution. Thus, uMRD cells exhibit distinct cell states from pre- and postSCT samples that comprise a wide spectrum of molecular phenotypes even within specific cell types.

### Clonal evolution following the SCT bottleneck

We found significant phenotypic evolution at postSCT relapse, a population dynamic that could result from either cell outgrowth (driven by subclonal competition) or cell type transition (driven by cell plasticity). To distinguish these two possibilities, we sought to track coordinated genotype-phenotype dynamics of MDS cells throughout preSCT, CR, and relapse timepoints. Specifically, we reconstructed the subclonal structure of recipient-derived cells at single-cell resolution in 7 patients with sampling at all 3 timepoints. We developed a clonal tracing approach that integrates CNV posterior probabilities inferred by Numbat (*45*) from the RNA layer with mitochondrial variant probabilities estimated by an Expectation–Maximization algorithm from the ATAC layer (Fig. S13A). We were able to successfully reconstruct phylogenetic, subclonal relationships in 6 of 7 patients (Fig. S13B). For example, in patient P01, an early defining mutation, 1887A>T, gave rise to the major malignant lineage (subclone 2), which subsequently diversified into two lineages (Fig. 5A-B). The first lineage includes subclones 3 and 4, which share mutations such as 10359A>T and 874G>C, with further divergence marked by 2814G>C. The other lineage contains subclones 5 and 6, defined by sequential acquisition of 9041A>G and 15897G>A.

**Fig. 5.**
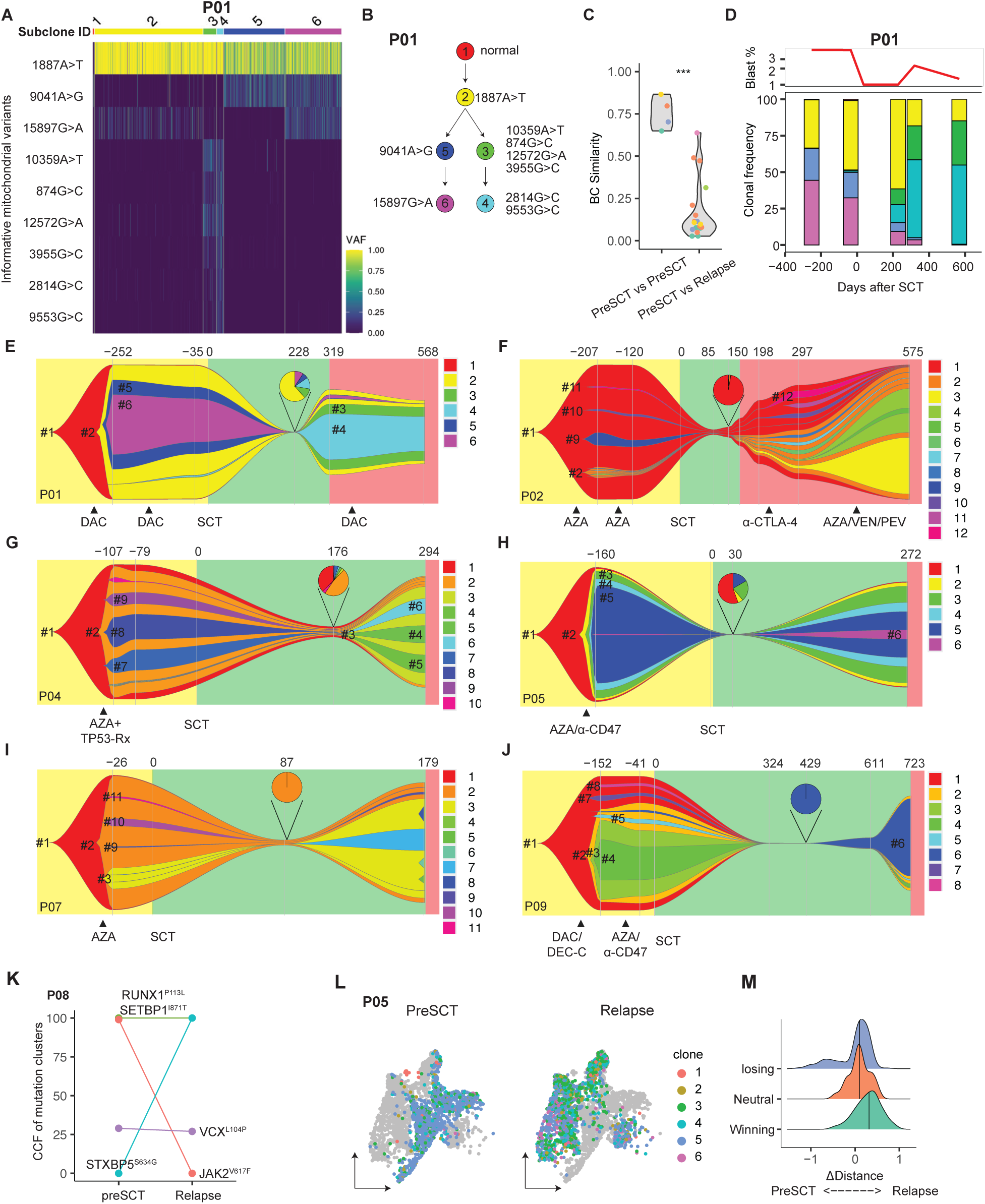
**Clonal dynamics and genetic changes from preSCT to relapse**. (A) Heatmap of mitochondrial variant allele frequencies used to infer clonal architecture across preSCT, CR, and relapse timepoints in patient P01. Colors indicate subclonal identity. (B) Phylogenetic tree showing subclonal relationships defined by shared mitochondrial mutations in patient P01. (C) Clonal composition similarity measured by Bray–Curtis index between sample pairs. Each point represents a pairwise comparison: “preSCT” compares one preSCT sample to another preSCT sample and “relapse” compares preSCT to relapse, all in the same patient. (D) Longitudinal clonal composition over time for patient P01; top panel shows blast cell percentage, bottom panel shows subclone fractions by day postSCT. (E–J) Fish plots of clonal evolution for patients with sufficient malignant cells at all timepoints, illustrating subclone dynamics from preSCT through CR (piechart) to relapse. (K) Changes in cancer cell fraction (CCF) of selected coding mutations identified from whole genome sequencing between preSCT and relapse for patient P08. (L) UMAP visualization of recipient-derived cells for patient P05, colored by clone assignment, showing state transitions for multiple subclones between preSCT and relapse samples. (M) Density plots of deviation scores for CD34 uMRD cells labeled as winning, neutral, or losing clones, comparing the phenotypic similarity of individual uMRD cells to preSCT versus relapse states. Lines for medians are shown for each group. Positive values indicate greater similarity to relapse, while negative values indicate greater similarity to preSCT. DAC, Decitabine; AZA, Azacitidine; α-CTLA-4, ipilimumab; VEN, Venetoclax; PEN, Pevonedistat; TP53-Rx, APR246; α-CD47, ALX148. DEC-C, Decitabine/cedazuridine.

As we did with <u>cell type</u> compositional similarity scores (Fig. S12B), we analyzed the <u>subclonal</u> compositional similarity scores across multiple pre- and postSCT relapse samples. As with the phenotypic shifts we measured (Fig. 4A), we also found significant changes in clonal composition after SCT across all patients (Fig. 5C) with distinct patient-specific patterns of subclonal structure and dynamics within the CD34 compartment (Fig. 5E-J). Nonlinear evolutionary dynamics were observed in 5 patients, in whom dominant subclones preSCT were replaced by subclones from distinct lineages after transplant. In patient P01, subclones #2, #5, and #6 were dominant prior to transplantation, with a frequency exceeding 50%; but at CR, #5 and #6 were nearly eradicated (falling below 5%) and replaced by subclones #3 and #4 at relapse (Fig. 5D-E). In patient P02, most of the cells prior to SCT belonged to subclone #1, which lacked mitochondrial mutations and CNVs, with the presence of a minor set of subclones (#3–8), all derived from subclone #2 and carrying a chr7 deletion (Fig. 5F and S13B). After SCT, subclone #1 remained dominant at uMRD and early relapse timepoints, but subclones #3-8 gradually expanded and became dominant at time of near-progression to AML with 18% marrow blasts. In patient P04, the preSCT population was dominated by subclone #2 and its descendants (#7–10), each with different mitochondrial mutations (Fig. 5G and S13C). After SCT, subclone #2 remained dominant at CR but was rapidly replaced within 4 months by subclones #3–6, characterized by chr1 and chr3 deletions, leading to relapse with sAML. In patient P05, subclones #2–6 evolved linearly and persisted both before and after transplantation without substantial subclonal substitution (Fig. 5H and S13D). In patient P07, the major pre-SCT population was subclone #2, characterized by chr3, 5, and 9 deletions, together with its descendants (#9–11) (Fig. 5I and S13E). Subclone #2 remained dominant at CR after transplant but was subsequently replaced at relapse by subclone #3. In patient P09, subclone #3, characterized by a chr5 deletion, and its descendants, were dominant before transplant but became very minor at CR and relapse (Fig. 5J and S13F), consistent with the clinical course in which the dominant MDS cells after transplant lacked CNVs. In summary, we observed significant subclonal sweeping in 5 of 6 patients, with clonal replacement occurring within a three-month sampling interval in most cases. In patients P01, P02, P04, and P07, minor subclones at CR expanded to become dominant at relapse, indicating that subclonal competition and evolution can occur during MRD progression in the absence of intervening therapy. In patient P09, the relapse-dominant subclone was already detectable at CR, suggesting that it either possessed increased resistance to SCT conditioning or had undergone clonal selection prior to uMRD sampling.

To confirm our observation of clonal replacement inferred from CARAMEL-seq, we further measured changes in coding, somatic single-nucleotide variants (sSNVs). We used whole-genome sequencing on bone marrow aspirate slide samples available from 6 patients (P01, P02, P06, P07, P08, and P09) at both preSCT and relapse. We tracked changes in the frequencies of identified somatic mutations (Fig. 5J and S14). Notably, all 6 patients exhibited mutations with greater than a 50% change in cancer cell fraction (CCF) between the two timepoints. For example, in patient P08, the CCF of the *JAK2* V617F mutation decreased from 99% preSCT to 0% postSCT (Fig. 5K). In patient P02, the CCF of *SETBP1* mutation decreased from 72% to 0%, while the CCF of *PTPN11* mutation increased from 20% to 99% between preSCT and relapse. These results substantiate that the clonal shifts we detected at single-cell resolution in the CD34 compartment are coupled with genetic changes representing the MDS bulk, in which CD34 cells constitute the majority. Thus, these data highlight the role of SCT in selectively eliminating dominant disease clones to shape MRD progression.

To determine whether the observed cell-state changes could be explained solely by clonal shifts, we next integrated clonal (genotype) with cell-state (phenotype) information for analysis. We found that cells from identical subclones occupied distinct regions in preSCT versus relapse samples (Fig. 5L), indicating that individual subclones undergo phenotypic reprogramming over time, in addition to shifts in their relative abundance. Moreover, this data suggests that phenotypic remodeling can occur in the absence of subclonal competition during MRD progression, as in P05.

We previously observed that in some patients, uMRD cells phenotypically resembled preSCT states, whereas in others they more closely resembled relapse states (Fig. 4E). We therefore asked whether this divergence is influenced by the clonal identity of uMRD cells. We incorporated clone-type labels and compared the phenotypic similarity of individual uMRD cells to relapse and preSCT cells. We defined strongly expanding clones with log fold change ≥ 0.5 at relapse as “winning” clones, weakly expanding clones with log fold change ≤ –0.5 as “losing” clones, and the remainder as “neutral.” We found that cells from winning clones exhibited higher phenotypic similarity to relapse than preSCT (Fig. 5M). The strong preSCT similarity bias observed in P07 and P04 within HSC/MPPs and LMPP-like cells may be explained by their origin in losing clones (Fig. S15). However, we also identified losing clones whose cells still displayed greater similarity to relapse (Fig. S15), indicating that subclonal identity alone is insufficient to explain phenotypic shifts in uMRD and relapse cells. Thus, phenotypic fates are heavily influenced by, but not exclusively bound to, subclonal identity. Indeed, the variation in phenotypic range across all subclones indicates the potential role plasticity may play in the survival of any given subclone. Taken together, these results suggest that SCT drives significant subclonal changes, with most patients exhibiting clear shifts during MRD progression. Furthermore, the data demonstrate that both subclonal competition and cell plasticity/state transitions shape MRD progression to drive the phenotypic remodeling seen at relapse postSCT.

### Increased malignant HSPC cytokine signaling and molecular remodeling during MRD progression

Given that MRD progression involved extensive phenotypic state transitions from preSCT to relapse across patients, we next aimed to identify potential molecular drivers of relapse. Since prior studies have shown that downregulation of HLA-class II proteins after SCT in AML is a common driver of relapse (*46, 47*), enabling resistance to T-cell attack, we first examined HLA-class II changes after relapse. We specifically analyzed the HSC/MPP-like cells and found that two of the 7 patients exhibited further reduction in HLA class II surface protein expression after SCT, suggesting a similar but uncommon phenomenon in our cohort (Fig. S16A). However, we found that most patients already exhibited lower HLA-class II expression at preSCT compared to donor baseline (Fig. S16B, table S1), indicating that impaired antigen presentation is an early property of MDS HSC/MPP-like clones before SCT rather than solely acquired at relapse post-SCT.

Next, we focused the altered gene expression and pathway activities shared among patients. Pseudobulk differentially expressed gene (DEG) analysis of HSC/MPPs comparing relapse to preSCT samples revealed that, despite the genetic and phenotypic heterogeneity of our cohort, several DEGs were consistently identified across patients (Fig. 6A). The most upregulated gene was *PYHIN1*, which encodes an interferon-inducible protein (*48*) and positively regulates IL-6 and TNF-α production (*49*). In contrast, downregulated genes in relapse included *ACSM3* and *SLC25A21,* both encoding mitochondrial proteins involved in acyl-CoA metabolism and reported as tumor suppressors in AML (*50–52*). Consistent with alterations in both inflammatory pathway and mitochondrial activities in HSC/MPPs after relapse, GSEA analysis revealed the enrichment of IFN-γ, TNF-α, IL-6, TGF-β, Notch, and KRAS pathway genes, while MYC, E2F, and cell cycle–related pathway genes were underrepresented in relapse (Fig. 6B). Indeed, cell-cycle phase inference indicated that fewer cells in relapse samples were actively cycling (Fig. 6C). These results highlight the global increase in inflammatory cytokine activities and decrease in cell division in HSC/MPPs at relapse.

**Fig. 6.**
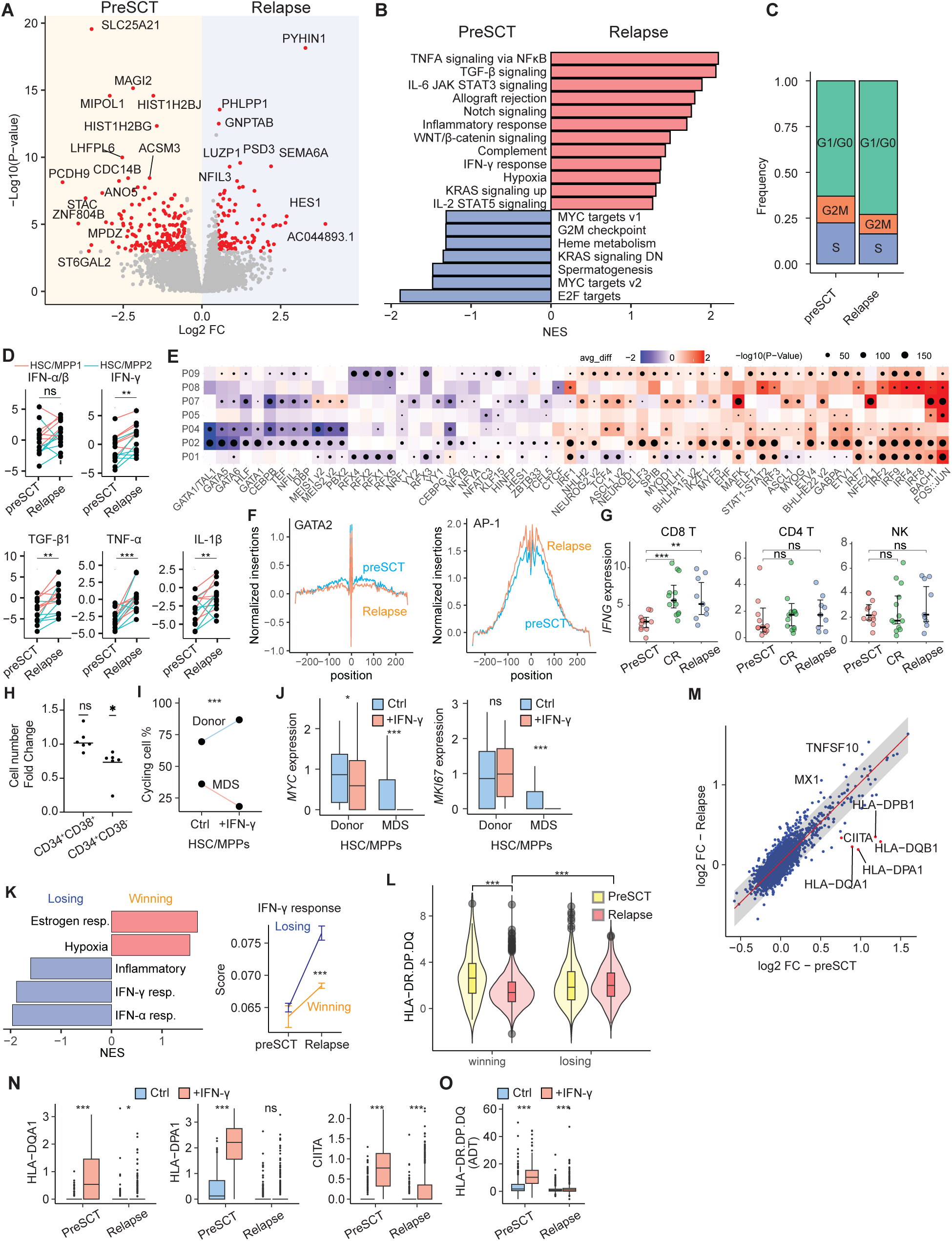
Transcriptional and functional remodeling driven by inflammatory cytokine signaling from preSCT to relapse. (A) Volcano plot of differentially expressed gene (DEG) analysis between relapse and preSCT HSC/MPPs using pseudobulk method; positive values indicate overexpression in relapse, with p-values floored at 10^-20^. (B) GSEA analysis of DEG results, where positive NES indicates overrepresentation in relapse and negative NES indicates overrepresentation in preSCT. (C) Cell-cycle phase ratios of HSC/MPPs from relapse and preSCT in single-cell analysis. (D) IFN-α/β, IFN-γ, IL-1β, TGF-β1, and TNF-α signaling activities inferred by CytoSig in HSC/MPPs from preSCT and relapse; each line represents a sample, with cells from HSC/MPP1 (red) or HSC/MPP2 (blue) clusters. p-values were determined by Wilcoxon test. (E) Differential eRegulon activity between relapse and preSCT HSC/MPP cells across patients. Heatmap shows only TFs with significant differences (FDR < 0.05), with red indicating higher activity at relapse. (F) TF footprinting profiles at GATA2 and AP-1 motifs. (G) Normalized IFNG expression in CD8 T cells, CD4 T cells, and NK cells across preSCT, CR, and relapse samples using pseudobulk approach. (H) Cell number fold change following IFN-γ treatment versus control in CD38 and CD38 compartments; p-values are based on H : fold change = 1 using t-test, with cell numbers measured by CellTiter-Glo® 2.0 Cell Viability Assay. (I) Cell-cycle phase inference from single-cell analysis after treating patient P09 HSC/MPPs with IFN-γ. (J) *MYC* and *MKI67* RNA expression in donor- versus recipient- derived HSC/MPPs under control and IFN-γ treatment. (K) Gene set enrichment and pathway activity analysis of winning and losing clones in patient P07 at relapse. Left, normalized enrichment scores (NES) of pathways with adjusted *p*-value < 0.05. Right, IFN-γ response scores in winning and losing clones across preSCT and relapse. (L) Violin plots showing HLA class II protein expression in winning and losing clones at preSCT and relapse. (M) Differential gene fold-change analysis after IFN-γ treatment. Each dot represents a gene. Scatterplot shows log fold change (logFC) in relapse (y-axis) versus preSCT (x-axis) samples. (N) HLA-DQ/DP and *CIITA* RNA expression and HLA class II protein expression (O) under control and IFN-γ treatment in patient P09 samples from preSCT and relapse. ns, not significant, *P < 0.05, **P < 0.01, and ***P < 0.001.

We next investigated which cytokine signaling pathways were consistently upregulated across patients. We performed the analysis by aggregating cells based on patient, cell type, and sample, followed by CytoSig to infer cytokine activities for the pseudobulked cells. The analysis revealed upregulation of IFN-γ, IL-1β, TNF-α, and TGF-β activity in relapse across patients but not type I IFN signaling (Fig. 6D and S17A).

Inflammatory cytokines can induce alterations in various transcription factor (TF) activities. To identify which TF activities were altered after relapse, we compared the activities of regulons, defined as a TF together with its predicted target genes, between HSC/MPPs from relapse and pre-SCT samples. Leveraging joint scRNA-seq and scATAC-seq profiles, we *de novo* reconstructed 333 enhancer-driven regulons (eRegulons) for 116 TFs with SCENIC+ (*53*). The analysis revealed the increased activities of KLF13, KLF1, NF-κB family members, and interferon regulatory factor 1 (IRF1), a key mediator of IFN-γ–induced signaling (Fig. S17B) and IRF4. In contrast, downregulated regulons after relapse included GATA1 and GATA2, known to be inhibited by IFN-γ (*54, 55*). Next, we analyzed TF-associated accessibility and performed motif enrichment of differentially accessible peaks as complementary approaches to infer changes in TF activity in HSC/MPPs. Consistent with regulon analysis, we detected enriched motifs for AP-1 (FOS:JUN), NF-κB, and multiple interferon regulatory factors (IRF1, 2, 4, 7, 8, and 9) at relapse and enriched GATA1/2 motifs at preSCT timepoints (Fig. 6E and S17C). The ATAC-based TF footprinting analysis further supported the increased activities of AP-1 and decreased activities of GATA2 (Fig. 6F). Thus, MDS relapse postSCT is characterized by heightened inflammatory signaling (e.g. IFN-γ and TNF-α) in malignant HSC/MPPs, evident in both transcriptional profiles and chromatin accessibility.

These signaling changes may stem from extrinsic factors, such as elevated cytokine levels in the bone marrow microenvironment, and intrinsic factors, such as altered receptor expression and signal transducer activity. Supporting an extrinsic source, we observed increased *IFNG* expression in postSCT CD8 T cells (Fig. 6G), but not in postSCT NK or CD4^+^ T cells, suggesting elevated IFN-γ secretion from CD8^+^ T cells. By contrast, the expression of IFN-γ receptor genes (*IFNGR1* and *IFNGR2*) did not show a significant increase in malignant HSC/MPPs at relapse (Fig. S17D). Thus, relapse-associated, cytokine signaling changes may be mainly driven by CD8^+^ T cell-derived IFN-γ secretion, though potential contributions from intrinsic mechanisms cannot be excluded.

We next evaluated the direct effects of IFN-γ on MDS cells. Since IFN-γ signaling upregulation coincided with downregulation of MYC pathway and cell cycle (Fig. 3E and 6B), and the suppression of *MYC* by IFN-γ has been reported in other cell types (*38*), we tested whether IFN-γ reduces cell proliferation and downregulates *MYC* in MDS cells. *Ex vivo* cultures of MDS cells from preSCT and relapse samples of patients P01, P02, and P09 revealed that IFN-γ reduced CD34 CD38 but not CD38 cell numbers after six days of expansion (Fig. 6H), indicating a cytostatic activity specific to the HSC/MPPs. Single-cell RNA-seq analysis of IFN-γ–treated samples from patient P09 further revealed that IFN-γ decreased the proportion of cycling cells in the MDS compartment while increasing it in the donor cell HSC/MPPs (Fig. 6I). Consistently, IFN-γ decreased both *MYC* and *MKI67* expression in MDS cells (Fig. 6J). These results indicate that IFN-γ promotes strikingly disparate fates in normal and malignant HSC/MPPs with decreased cell cycle and MYC activity in the latter.

We then took two approaches to glean insights into the possible role of IFN-γ signaling as a stressor during MRD progression. First, we compared pathway activity between the winning and losing subclones at relapse across the patients. While all patients demonstrated increased IFN-γ signaling at relapse (Fig 6B, D), discrepant IFN-γ signaling between winning and losing subclones was detected only in P07, possibly indicating different roles for this pathway in driving divergent evolutionary trajectories across patients (Fig. 6K). Consistent with the decreased IFN-γ signaling in the winning subclones in P07, we also detected decreased RNA and surface protein expression of known, IFN-γ-inducible HLA class II molecules in winning subclones (Fig. 6L). HLA class II molecules are known activators of potent graft-versus-leukemia (GvL) effects, and their downregulation on the winning subclone suggests that, in P07, heightened IFN-γ signaling may contribute to selective pressure against susceptible subclones; and that the winning clone might attenuate IFN-γ to avoid HLA class II upregulation and evade anti-leukemic T cell immunity.

Second, we functionally examined differential responses of matched, preSCT and relapse MDS cells to IFN-γ perturbation *in vitro*. Because P07 relapsed fewer than 6 months (179 days) after SCT, we studied P09 who relapsed ∼24 months (723 days) after SCT (the longest time to relapse in our cohort). Single-cell analysis of recipient-derived HSC/MPP-like MDS cells revealed that gene-level fold changes by IFN-γ between preSCT and relapse were highly correlated (ρ=0.77). We next fit these fold changes using a linear regression model with the slope constrained to 1, which showed that 99.8% of genes deviated by less than ±0.37 log FC from the fitted line, indicating broadly preserved IFN-γ responses across the two timepoints. Strikingly, of the 0.2% of genes whose IFN-γ regulation was not conserved, seven with the largest residuals included HLA class II genes (*HLA-DQA1*, *HLA-DQB1*, *HLA-DPA1*, *HLA-DPB1*) and their key regulator, *CIITA* (Fig. 6M). Both genes and corresponding surface proteins were significantly less inducible by IFN-γ in relapse compared to preSCT cells (Fig. 6N and O) with low baseline expression levels at both timepoints, suggesting a rewiring specifically of and limited to IFN-γ– induced HLA class II expression. Altogether, the *in-silico* measurements and *in vitro* perturbation experiments indicate that IFN-γ may serve as a selective stressor after SCT, favoring the subclones that can achieve specific resistance to IFN-γ–induced HLA class II expression along distinct paths, potentially enabling immune evasion and contributing to disease persistence and relapse.

## Discussion

MRD has been widely recognized as a harbinger of relapse in MDS and AML; yet its biological nature and evolutionary trajectory have remained poorly understood. In this work, we sought to develop sensitive tools to comprehensively profile MRD and provide a mechanistic framework for how residual clones persist and evolve under therapeutic and immune pressure. We devised a highly sensitive assay that surpasses multiparameter flow cytometry methods and resolves the subclonal composition, transcriptional programs, and immunophenotypes of MRD cells. Applied to longitudinal sampling, we revealed the molecular dynamics of MRD progression after SCT.

The strength of CARAMEL-seq lies in its ability to integrate multiple orthogonal features to resolve MRD with high confidence, in contrast to existing methods that rely on single-layer information (*13–16, 56*). By leveraging tools such as Souporcell, we can demultiplex donor- and recipient-derived cells and compare residual recipient populations against donor-derived RNA, protein, and mitochondrial variant profiles that serve as a robust baseline. This multimodal framework is particularly powerful in MDS, a clinically heterogeneous disease in which patients harbor distinct immunophenotypic alterations: some exhibit strong surface-marker shifts, whereas others show minimal immunophenotypic change but may carry complex cytogenetic abnormalities (*57, 58*). For cases lacking stable phenotypic markers or cytogenetic abnormalities, mitochondrial variants provide an additional independent layer for lineage tracing and subclonal tracking. The integration of these diverse modalities also ensures that residual clones can be identified reliably even as their defining features evolve. For example, in one patient (P09), the dominant preSCT clone carried a del(5q), yet after transplant, the dominant residual population was cytogenetically normal; our method nevertheless tracked these cells across disease states. Thus, the combination of donor–recipient demultiplexing, multimodal baselines, and redundant molecular markers overcomes the limitations of single-modality assays and provides a comprehensive platform for MRD detection.

Applying this framework, we uncovered several key features of MRD biology (Fig. 7). In contrast to the prevailing view that therapeutic bottlenecks inevitably reduce clonal diversity (*59, 60*), our data show that MRD retains substantial intrapatient heterogeneity even after the intense selective pressure of conditioning regimens. Residual disease persists as genetically and phenotypically distinct subclones, yet within this complexity, we identified shared transcriptomic and surface proteomic signatures across patients. The shared higher expression of *PTH2R*, CD244, and GPR56 genes and proteins, respectively, presents potential new MRD markers and directs future mechanistic interrogation to understand their functional relevance to MRD.

**Fig. 7.**
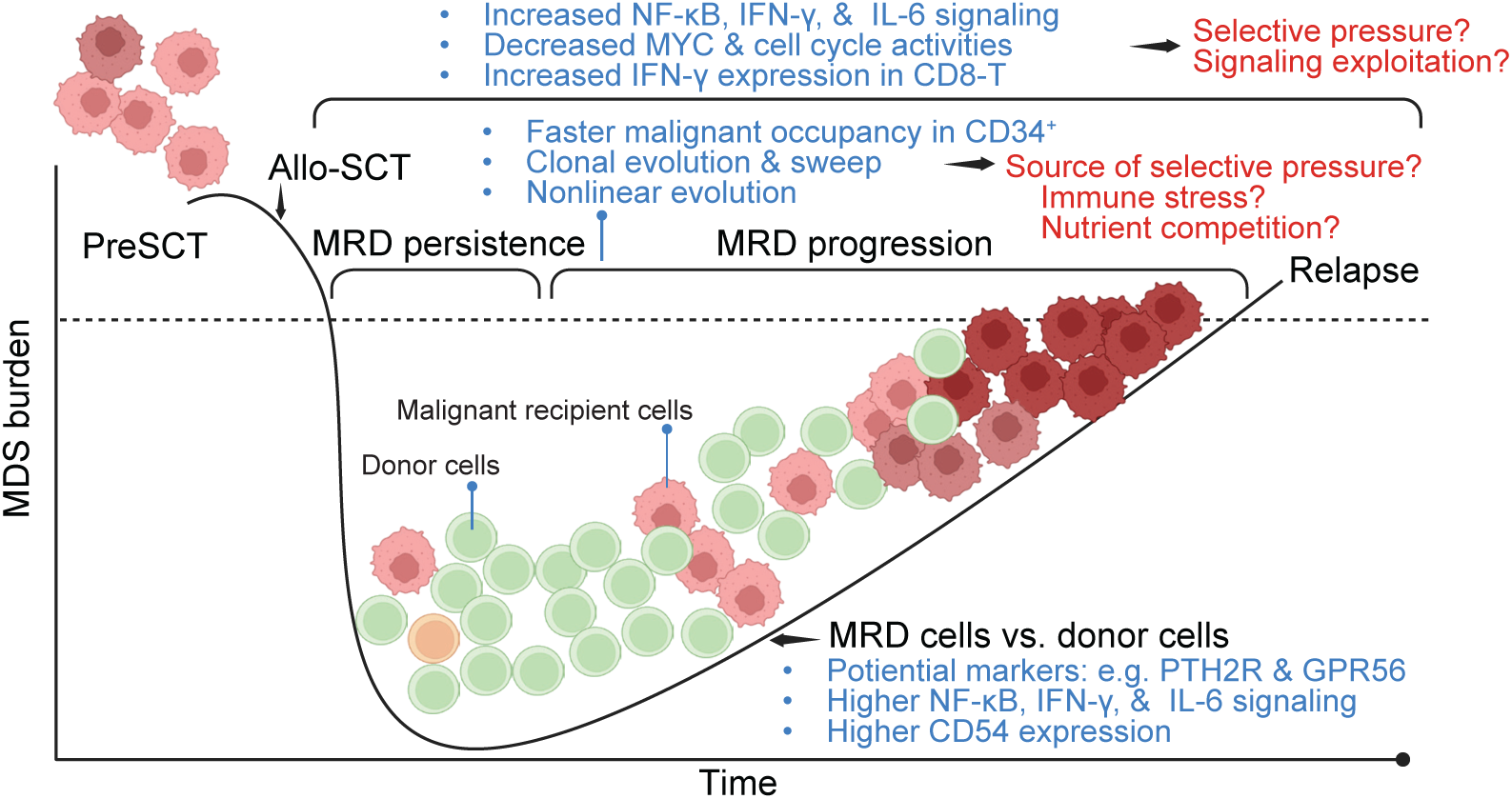
**Clonal and molecular dynamics of MDS MRD after SCT**. Schematic of disease burden from preSCT to relapse. Malignant cells (red) showed faster occupancy of CD34 compartments than CD34 . MRD progression was characterized by clonal evolution and sweeping, nonlinear evolution. Key molecular features after transplant included increased NF-κB, IFN-γ, and IL-6 signaling as well as reduced MYC and cell-cycle activity in malignant HSCs; and higher IFN-γ expression in CD8^+^ T cells. Compared to donor cells, MRD shows increased expression of *PTH2R* and GPR56, NF-κB/IFN-γ/IL-6 signaling, and CD54 expression.

Longitudinal sampling revealed that MRD progression is dynamic rather than static or merely expansionary. By capturing samples at CR and at the earliest stages of relapse, an interval without intervening or maintenance therapy in our cohort, our workflow enabled reconstruction of the natural history of MRD evolution. Coupled with our single-cell multi-omic workflow and new tool for subclonal inference, this approach allowed us to directly address the question of subclonal outgrowth versus state transition, which cannot be resolved by bulk RNA-seq or without clonal tracing. We found that phenotypic evolution between CR and relapse is common, and residual subclones of MRD differ markedly in their outgrowth potentials, with some extinguishing and others emerging as dominant drivers of relapse. These findings reposition MRD as an active evolutionary reservoir rather than a passive remnant and provide a framework for understanding how relapse trajectories are sculpted by both intrinsic clonal fitness and extrinsic selective pressures.

Mechanistically, our multi-layered analyses converged on cytokine signaling within malignant HSC/MPPs as a dominant signature after SCT, conserved across patients. We found that malignant HSC/MPPs exhibit heightened TNF-α, IFN**-**γ, IL-6, TGF-β and IL-1β signaling. Supporting a role for extrinsic drivers, we observed increased expression of *IFNG* in CD8^+^ T cells after SCT, indicating that SCT reshapes the immune microenvironment and the interplay between malignant stem/progenitor cells and surrounding immune populations. Co-occurring transcriptional, epigenetic, and clonal dynamics collectively suggest that these heightened signals can act as evolutionary pressures. Indeed, in one patient, the winning and losing subclones displayed differential IFN**-**γ signaling and concomitant HLA class II expression, indicating that cytokine exposure prunes residual diversity, selectively favoring subclones with adaptive fitness. These findings reinforce the importance of HLA class II downregulation in myeloid malignancies, consistent with prior reports showing that HLA class II decreases after SCT and that such downregulation impairs CD4 T cell–mediated GvL activity in AML (*46, 47*). However, unlike these earlier observations, we found that HLA class II downregulation relative to normal CD34^+^ is already common prior to SCT in our MDS cohort, suggesting that impaired antigen presentation occurs before SCT. Moreover, in another patient with late relapse, cytokine perturbation experiments revealed that subclones had rewired their IFN-γ responses by acquiring a specific defect in HLA class II inducibility, highlighting an additional layer of immunoediting. Nevertheless, we acknowledge and indeed expect that evolutionary trajectories during MRD progression may differ across patients. Future large-scale studies will be required to delineate the distinct types of clonal adaptations arising under the common cytokine signaling changes after SCT, thereby enabling patient stratification and the development of personalized treatment strategies. In summary, these findings recast MRD as a genetically diverse but phenotypically convergent evolutionary reservoir, shaped by immune and therapeutic pressure, and identify cytokine-driven maladaptive signaling as a tractable axis to intercept relapse.

The multimodal framework established by CARAMEL-seq opens several avenues. Translationally, MRD-selective markers identified in this study could be incorporated into next-generation flow cytometry or single-cell diagnostic assays to improve clinical monitoring and inform postSCT maintenance therapy strategies. Fundamentally, the eco-evolutionary forces sculpting MRD suggest that dissecting the interplay between cytokine-driven signaling, immune editing, and clonal fitness will be essential for understanding why certain subclones escape control while others succumb. Larger patient cohorts and longitudinal sampling will be required to validate the generality of these mechanisms across MDS and related malignancies. Therapeutically, strategies that disrupt maladaptive cytokine responses—particularly IFN-γ signaling—should be explored to alter the evolutionary trajectory of MRD progression and prevent relapse. Ultimately, such molecular toolkits could be broadly applied across cancer to redefine MRD from a poor prognostic sign to an effective therapeutic target.

## Acknowledgments

We thank Guillermo Montalban Bravo, Danielle Hammond, Kelly Chien, Simona Colla and all members of the Bachireddy laboratory for valuable discussions. We appreciate faculty members from the Department of Hematopoietic Biology & Malignancy for their helpful feedback on the manuscript. Library sequencing was performed by The Advanced Technology Genomics Core, supported in part by The University of Texas MD Anderson Cancer Center and P30CA016672. Flow cytometry experiments were performed in and assisted by The Advanced Cytometry & Sorting Facility at South Campus (ACSF). We are grateful for the study nurses and clinical staff that obtained samples, the research staff who processed and banked these samples, and the patients who generously consented for the research use of these samples. This work was supported in part by the PhRMA foundation (to Y.-H.C), Edward P. Evans Foundation (to P.B.), Damon Runyon Cancer Research Foundation (to P.B.), Cancer Prevention and Research Institute of Texas (RR210008 to P.B.), the Break Through Cancer Foundation (to P.B.), and NCI grant 1K08CA248458-01 (to P.B.). P.B. is a CPRIT Scholar in Cancer Research and an Andrew Sabin Family Foundation Fellow at The University of Texas MD Anderson Cancer Center. A.M.V. received funding from the ASH-CIBMTR-ASTCT Career Development Award for this project.

## Author contributions

Y.-H.C. and P.B. conceived the study. Y.-H.C., A.P., and P.B. interpreted the results of the data and supervised the study. Y.-H.C. designed and performed experiments. Y.-H.C., T.B., M.X., and M.Z., analyzed the results. H. Y. collected and processed BM samples from patients. G. A.-A., G.R., E. S., J.M., D.M., S.G., and G.G.-M. oversaw patient care and provided clinical data from the BMT repository. A.M.V. and F.C.-C. organized the clinical data. F.J., C.K., P.K. J.L., and W.L. assisted and performed whole-genome sequencing. Y.-H.C., T.B., M.X., and P.B. wrote the manuscript with input from all authors.

Diversity, equity, ethics, and inclusion [optional]:

## Data and materials availability

The sequencing data generated in this study have been deposited in the NCBI BioProject database under accession number PRJNA1302443.

The analysis scripts used to reproduce results included in the paper are available at https://github.com/yuhsiangchen-bioinfo/MDS-MRD-project.

## Supplementary Materials

### Materials and Methods

#### Bone marrow samples

Bone marrow aspirates from patients with MDS were collected at the University of Texas MD Anderson Cancer Center (Houston, TX) following written informed consent under MDACC Institutional Review Board-approved protocols according to the principle of the Declaration of Helsinki and the International Conference on Harmonization Guideline for Good Clinical Practice. Diagnoses were confirmed by expert hematopathologists. Bone marrow mononuclear cells (BMMCs) were isolated and cryopreserved using Ficoll-Paque PLUS density gradient centrifugation (GE Healthcare).

#### Sample processing

Cryopreserved BMMCs were thawed at 37 °C on the day of sequencing. Cells were immediately transferred to a new tube and gradually diluted with a pre-warmed thawing buffer consisting of 50% fetal bovine serum (FBS), 25% anticoagulant solution (50 mM sodium citrate, 25 mM citric acid, 82 mM dextrose), and 25% 0.9% sodium chloride (NaCl). Thawing buffer was added in three sequential 1:1 volume increment at a rate of 1 mL per 3 seconds while gently swirling. Cells were centrifuged at 300 × g for 8 minutes at room temperature, and the supernatant was carefully aspirated, leaving ∼1 mL without resuspension. Nine milliliters of media were then added dropwise at the same rate to reach a final volume of 10 mL, followed by centrifugation at 300 × g for 5 minutes. The supernatant was again removed, leaving ∼0.5 mL, and 5 mL of MACS buffer was gently added without disturbing the pellet. A final centrifugation was performed at 300 × g for 8 minutes at 4 °C. The supernatant was removed, and the cell pellet was resuspended in 100 µL of MACS buffer containing autoMACS® Rinsing Solution supplemented with 1% bovine serum albumin (BSA).

#### Microsatellite-based chimerism

Microsatellite-based testing for assessment of stem cell engraftment (chimerism) was performed in the University of Texas MD Anderson Cancer Center Molecular Diagnostic Lab on genomic DNA extracted from peripheral blood or bone marrow aspirate samples. For peripheral blood, additional testing was conducted on DNA isolated from sorted T lymphocytes, myeloid cells, and/or NK cells, when applicable.

#### Somatic mutation testing of leukemia driver genes

Genomic profiling of recurrent leukemia driver genes was performed in the Molecular Diagnostic Laboratory, The University of Texas MD Anderson Cancer Center, using a hybridization capture–based NGS platform. Sequencing libraries were prepared from genomic DNA isolated from fresh bone marrow aspirates, peripheral blood, or formalin-fixed paraffin-embedded (FFPE) bone marrow clot specimens. DNA was subjected to target enrichment with the HaloPlex™ system (Agilent Technologies, Santa Clara, CA), capturing either entire coding sequences or recurrent hotspots across 81 genes commonly mutated in hematologic malignancies (*61*). The panel incorporated molecular barcodes to improve sensitivity and reduce sequencing artifacts. Paired-end sequencing was performed on the Illumina MiSeq platform (Illumina, San Diego, CA).

#### Fluorescence activated cell sorting (FACS)

Up to 5 million cells were incubated with 5 µL of Fc receptor blocking reagent (BioLegend, cat. no. 422302) in 100 µL of MACS buffer for 5 minutes at 4 °C. Subsequently, cells were stained with 1 µL anti-CD34-PE (BioLegend, cat. no. 343504), 10 µL anti-CD45-APC (BD, cat. no. 340942), 1.5 µL of 0.5 mg/mL TotalseqA Hashtag antibody, 0.5 µL anti-CD34-TotalseqA (BioLegend, cat. no. 343537), and 0.5 µL anti-CD45-TotalseqA (BioLegend, cat. no. 304064). Staining was performed on ice for 30 minutes in the dark. Cells were then washed twice with 1 mL of MACS buffer by centrifugation at 300 × g for 8 minutes at 4 °C, and supernatants were carefully aspirated. Finally, cells were resuspended in 0.5 mL of MACS buffer containing 1.5 µL of 1 mg/mL 7-AAD viability dye (Invitrogen™, cat. no. A1310). Cells were sorted on a BD FACSAria™ sorter into two populations: (1) CD34 and (2) CD34 CD45 cells —using Eppendorf tubes containing 20 µL of Cell Staining Buffer (BioLegend, cat. no. 420201) as collection tubes.

#### Cell staining with barcoded antibodies

Sorted CD34 and CD34 CD45 cells from different timepoint samples were combined and transferred to a 96-well plate (SARSTEDT, cat. no. 83.3926.500) for staining with barcoded antibodies. TotalSeq™ antibody cocktail was prepared by reconstituting the TotalSeq™-A Human Universal Cocktail, V1.0 (BioLegend, cat. no. 399907) with 42 µL of staining buffer, followed by the addition of 1 µL each of the following TotalSeq™-A antibodies: anti-CD90 (BioLegend, cat. no. 328135), anti-CD10 (BioLegend, cat. no. 312231), anti-CD15 (BioLegend, cat. no. 323046), anti-CD117 (BioLegend, cat. no. 313241), anti-CD133 (BioLegend, cat. no. 394005), anti-MICA/MICB (BioLegend, cat. no. 320921), anti-CD34 (BioLegend, cat. no. 343537), and anti-CD45 (BioLegend, cat. no. 304064). 25 µL of the prepared antibody cocktail was added to each sample well. Cells were incubated for 30 minutes at 4 °C. Cells were then washed five times with 200 µL of staining buffer by centrifugation at 300 × g for 5 minutes at 4 °C. After the final wash, cell pellets were resuspended in 40 µL of DPBS with 0.04% BSA for downstream processing.

#### Cell fixation and permeabilization for DOGMA-Seq/TEA-Seq

DOGMA/TEA-seq workflows were used to obtain single-cell RNA, ATAC, and protein profiles (*62*, *63*). To fix cells, 4.6 µL of 4% formaldehyde (Thermo Scientific Chemicals, cat. no. J60401.AK) was added (final conc. 0.1%) and gently pipette-mixed three times. Cells were incubated at room temperature for 5 minutes. Fixation was quenched by adding 4.4 µL of 1.25 M glycine (final conc. 0.125%) with gentle mixing. Cells were then washed twice with 160 µL of ice-cold PBS/BSA/RI buffer—comprising 1% BSA and 0.2 U/µL RNase inhibitor (Roche, cat. no. 3335402001) in DPBS—and centrifuged for 5 minutes at 400 × g at 4 °C.

The resulting cell pellet was permeabilized in 25 µL of chilled LLL lysis buffer, consisting of 10 mM NaCl, 3 mM MgCl, 1% BSA, 1 mM DTT, 0.1% IGEPAL CA-630 (Thermo Scientific, cat. no. J19628-AP), and 2 U/µL RNase inhibitor in nuclease-free water and gently mixed three times. Cells were incubated on ice for 3 minutes. Subsequently, 225 µL of chilled LLL wash buffer—composed of 10 mM NaCl, 3 mM MgCl, 1% BSA, and 1 mM DTT in nuclease-free water—was added, and the mixture was gently mixed and transferred to a new well. Cells were centrifuged for 5 minutes at 500 × g at 4 °C. A final wash was performed with 250 µL of LLL wash buffer, and the cells were resuspended in 5 µL of 1× diluted nuclei buffer (10X Genomics, part no. 2000207). The suspension was carefully pipetted and counted, and the cell concentration was subsequently adjusted according to 10X Genomics single-cell assay loading guidelines.

#### Transposition and barcoding for DOGMA-Seq/TEA-Seq

Cells were processed according to the Chromium Next GEM Single Cell Multiome ATAC + Gene Expression User Guide (10x Genomics, CG000338 Rev D), with several modifications. During the pre-amplification PCR step, 1 µL of 0.2 µM ADT additive primer (5’ CCT TGG CAC CCG AGA ATT CC) and 1 µL of 0.2 µM HTO additive primer (5’ GTG ACT GGA GTT CAG ACG TGT GCT CTT CCG AT*C* T, *a phosphonothioate bond) were added to the reaction mix. Following pre-amplification PCR and SPRI cleanup, beads were eluted in 100 µL of EB buffer. From the eluate, 25 µL was used for ATAC library indexing and 30 µL was used for cDNA amplification. To amplify HTO and ADT libraries, two separate PCR reactions were performed using pre-amplified material: 25 µL for ADT and 5 µL for HTO. Each reaction included the SI-PCR primer (5′-AAT GAT ACG GCG ACC ACC GAG ATC TAC ACT CTT TCC CTA CAC GAC GC*T* C) along with RPI-x indexing primers for ADT and D70x_Long primers for HTO, each at a final concentration of 0.25 µM. PCR was carried out in a total volume of 100 µL using 50 µL of 2× KAPA HiFi HotStart ReadyMix (Roche cat. no. 07958927001). Thermocycling conditions were: 95 °C for 3 minutes, followed by 13–15 cycles of 95 °C for 20 seconds, 60 °C (ADT) or 64 °C (HTO) for 30 seconds, and 72 °C for 20 seconds; with a final extension at 72 °C for 5 minutes. Final libraries were quality-assessed using High Sensitivity DNA chips (Agilent, cat. no. 5067-4626) on a Bioanalyzer 2100 system.

#### MAESTER-Seq library preparation

For samples with limited CD34 cell counts—including complete remission samples and all patient P13 samples—the Chromium Next GEM Single Cell 3′ Reagent Kits v3.1 (Dual Index) were used according to the manufacturer’s protocol to avoid cell loss associated with cell permeabilization in DOGMA/TEA-seq workflows. As these samples lacked ATAC information to capture mitochondrial DNA, the MAESTER-seq protocol (64) was applied to recover single-cell mitochondrial variant profiles from RNA layer.

MAESTER-seq workflow includes a two-step PCR amplification. In the first PCR, 12 separate reactions were set up to amplify tiled regions across the mitochondrial transcriptome using a 12- primer mix. Each reaction contained 0.1 μM of each target-specific primer, 1 μM GoT-P5-i5- BCXX primer for indexing, and 1× KAPA HiFi HotStart ReadyMix in a final volume of 39 μL, with 1 μL of cDNA (from the 10X cDNA amplification step) as input. Thermocycling conditions were as follows: initial denaturation at 95 °C for 3 minutes; 8 cycles of 98 °C for 20 seconds, 65 °C for 15 seconds, and 72 °C for 3 minutes; followed by a final extension at 72 °C for 5 minutes.

Following the first amplification, PCR products were pooled in defined proportions, purified using 1× SPRIselect beads (Beckman Coulter, cat. no. B23318), and eluted in 20 μL of nuclease-free H O. For the second PCR to add P7 Illumina adapters, 18 μL of the eluate was combined with 20 μL of 2× KAPA HiFi HotStart ReadyMix, 1 μL of P5-generic primer (10 μM), and 1 μL of P7-i7-BCXX primer (10 μM). The PCR conditions were initial denaturation at 95 °C for 3 minutes; 6 cycles of 98 °C for 20 seconds, 60 °C for 30 seconds, and 72 °C for 3 minutes; followed by a final extension at 72 °C for 5 minutes. After the second PCR, the amplified DNA was purified using 0.8× SPRIselect beads and eluted in 20 μL of nuclease-free H O.

#### CD54 expression on MDS-L cell line with cytokine stimulation

MDS-L cells (*65*) were cultured in RPMI-1640 medium supplemented with 10% FBS, 1% penicillin-streptomycin, and 20 ng/mL IL-3. Cells were stimulated with 20 ng/mL human recombinant IFN-γ or 20 ng/mL TNF-α (STEMCELL Technologies, cat. no. 78068.1) with unstimulated cells served as controls. After 3 days, cells were washed and stained with anti- CD54-BUV737 (BD, cat. no. 741841). Flow cytometric analysis was performed using a BD LSRFortessa™ high-parameter flow cytometer.

#### Single-cell data processing

Raw sequencing FASTQ files from ATAC and RNA libraries were processed using CellRanger v7.0.0 for 10x Genomics 3′ kits v3.1 and CellRanger-ARC v2.0.1 for 10x Genomics multi-omics kits v1 with a nuclear-mitochondrial masked reference genome to improve mitochondrial read mapping. The masked reference genome was generated and indexed following the mgatk guidelines. For ADT and HTO libraries, the count matrices were generated from FASTQ files using the kite, kallisto, and bustools workflow (66–68).

RNA count matrices generated by CellRanger were processed using Seurat v4, and ATAC count matrices were processed using Signac (69). Low-quality cells were excluded based on the following criteria: fewer than 250 or more than 7,500 detected genes, fewer than 200 ATAC features, or over 15% mitochondrial gene content. The ADT counts were de-noised and normalized using dsb (70).

#### Sample- and genotype-based demultiplexing

Libraries containing HTO-multiplexed samples were demultiplexed using demuxmix (71) with a posterior acceptance threshold (pAcpt) of 0.6 based on HTO count data. To determine donor and recipient cell origin for individual cells based on genotypes, Souporcell was applied using the merged CellRanger BAM files along with the corresponding cell barcodes for each patient. To reduce misclassification, only cells with a singlet posterior probability equal to 1 in the Souporcell output were retained.

#### Cell-type annotation

A bone marrow single-cell multiomic CD34 reference map was constructed using donor-derived normal cells to enable cell-type label transfer on malignant cells for identifying their normal counterparts. First, RNA count matrices from donor cells were normalized and scaled using SCTransform (*72*), with mitochondrial percentage regressed out. PCA was performed on the SCTransform-corrected values using the top 30 dimensions. Batch correction was then applied on the PCA using Harmony (*73*), with sample identity as a covariate. For ATAC count matrices, term frequency–inverse document frequency (TF-IDF) normalization was first applied, followed by feature selection using FindTopFeatures function with a minimum cutoff at the 5th percentile (*69*). Dimensionality reduction was then performed using singular value decomposition (SVD), and the resulting dimensions were batch-corrected using Harmony. For ADT count matrices, dsb-normalized values were scaled and subjected to PCA, followed by batch correction. To integrate the three modalities, a weighted nearest neighbor (WNN) graph was constructed using batch-corrected dimensional reductions: components 1–30 from SCTransform (RNA), components 2–50 from ATAC, and components 1–30 from dsb-derived ADT data. Clustering and UMAP were performed on the weighted shared nearest neighbor (WSNN) graph using the SLM algorithm with the resolution parameter set to 1. Clusters were annotated as specific cell types based on marker gene and protein expression, as well as gene set enrichment analysis. Pseudotime trajectories were inferred using Monocle3.

Supervised PCA (SPCA), guided by a cell-cell kernel, was performed to identify an RNA-layer transformation that best captures the structure of the weighted nearest neighbor (WNN) graph. The resulting SPCA components were used as the reference dimensional reduction. Using Seurat v4 (74), anchors were identified between each recipient-derived query dataset and the multimodal reference, allowing recipient cells to be mapped and annotated through label transfer.

#### Multimodal UMAP and clustering with MultiVI

Patient-specific and atlas-level UMAPs were generated from the latent spaces learned by MultiVI. For atlas-level integration, all cells across samples were jointly modeled using RNA, ATAC, and protein modalities. Library technology (10X 3′ kit or multiome kit) and sample identity were included as batch covariates. The 23-dimensional latent space output from the model was exported and used to construct the UMAP embedding in Seurat v4.

For patient-specific UMAPs, MultiVI models (44) were trained separately for each patient by subsetting their respective cells. Experimental batch ID—where a single batch can include multiple HTO-labeled samples—was used as a batch covariate to correct for technical variation while preserving inter-sample biological differences. The 23-dimensional latent space outputs from the patient-specific MultiVI models were exported and used to construct UMAP embeddings in Seurat v4. Clusters derived from the latent space were interpreted as distinct cell states within each patient’s dataset.

#### Diversity and similarity metrics

Cell-type diversity

Cell-type diversity per donor or recipient compartment was quantified by Shannon entropy:

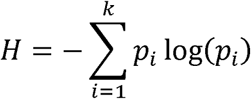

where pi is the proportion of cells belonging to cell type ; and k is the total number of cell types in the compartment.

Cell state and clonal similarity

For cell state and clonal composition similarity analysis, Bray–Curtis similarity scores were calculated across sample pairs (preSCT–preSCT and relapse–preSCT) to evaluate overall similarity in cell state or clonal composition between samples. Bray–Curtis similarity was defined as:

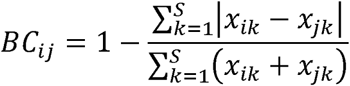

where xik and x}k denote the abundance of cell type (or subclone) k in samples, and,, and s is the total number of cell type (or subclone). Statistical significance was assessed using two-sided

Wilcoxon rank-sum tests.

Phenotypic shift

The phenotypic shift for each cell type c from preSCT to relapse was quantified using a dissimilarity index in the MultiVI latent space. For each cell type c, pairwise distances were measured within preSCT cells (Pc), within relapse cells (Rc), and between preSCT–relapse cells (vc). The dissimilarity index for cell type c was defined as:

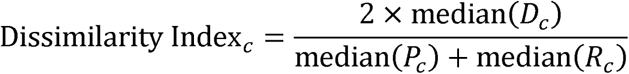

Similarity bias of MRD cells space from each MRD cell of cell type c to all preSCT cells of the same type (Pc1, …, Pci) and To quantify the phenotypic bias of MRD cells, we calculated distances in the MultiVI latent relapse cells of the same type (Rc1, …, Rc}). The ΔDistance for cell type c was defined as the difference between the median distance to relapse cells and the median distance to preSCT cells, capturing the relative displacement of MRD cells toward relapse versus preSCT identities.”

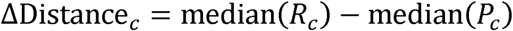

#### CNV calling from single cell data

Numbat v1.4.2 (45) was used to infer copy number variations (CNVs) on a per-patient basis from the scRNA-seq layer, executed within a Singularity container using the Numbat Docker image. BAM files from RNA libraries were merged and renamed by prefixing with the experiment ID and suffixing with the lane ID. SNP pileup data were generated using cellSNP-lite v1.2.3 (75) and phased using Eagle v2.4.1 (76) with the 1000 Genomes Project Reference Panel and SNP VCF, providing recipient cell barcodes for genotype assignment. Reference expression profile for CNV calling was generated from donor-derived cell RNA matrices aggregated by cell types.

#### Mitochondrial variants calling

For samples processed with the 10X multiomic kits, mitochondrial variant count matrices were derived from the ATAC library using mgatk v0.6.1 (77). Count matrices from each recipient were merged, and variants were identified using the IdentifyVariants function in Signac (69). Variants were retained if they were confidently detected in at least five cells (n_cells_conf_detected ≥ 5) and exhibited minimal strand bias (strand_correlation ≥ 0.65).

For samples processed with the 10X 3′ kits, mitochondrial variant count matrices were generated from the MAESTER-Seq library using maegatk v0.2.2 (64). FASTQ files were first trimmed using cutadapt and then aligned to the nuclear mitochondrial DNA segments (NUMT)-masked hg38 reference genome using STAR v2.7.9 (78). The resulting BAM files were processed with samtools, including the addition of cell barcode and UMI tags. Mitochondrial variants were quantified with maegatk using a minimum base quality of 20 for inclusion in genotype counts (-q 20) and a minimum of 3 supporting reads to call a consensus UMI/read (--min-reads 3).

#### Lineage tracing

Subclonal reconstruction and assignment were performed by integrating CNV and mitochondrial variant information using ScisTree (79), which uses single-cell genotype probabilities as input to deal with uncertainty. For mitochondrial genotypes, the observed variant counts in each cell depend on the total sequencing depth at each genomic position in each mutated cell and can be further influenced by cross-contamination of ambient fragments. To address these factors and provide genotype probabilities, we applied a negative binomial (NB) regression model t0 evaluate the relationship between mutant read counts (y) and mitochondrial base coverage (cov), incorporating a latent group variable (c) that distinguishes cells harboring mutant bases (c =1) from those without mutations (c = 0).

The mutant read counts (y) for mutation m were modeled as:

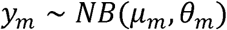

where µ is the mean parameter and 0 is the dispersion parameter of the Negative Binomial distribution for the mutation m. The mean parameter is defined as:

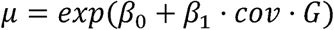

µ0 captures background mutant reads from cross-contamination or sequencing errors, representing baseline levels independent of mitochondrial base coverage or group assignment. µ1 quantifies the effect of mitochondrial base coverage (cov) in cells with the mutation m (c = 1). The latent variable c group takes binary values:

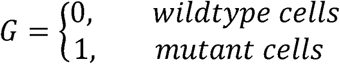

This formulation reflects the observation that mutant read counts do not scale with sequencing depth for cells without the mutation.

To estimate the parameters (µ0, µ1, 0) and infer the latent group variable (c), we employed the Expectation-Maximization (EM) algorithm.

Steps of the EM Algorithm:

1. Initialization: Model parameters were initialized as µ0 = − 2, µ1 = 1, and 0 = 2. The initial probabilities for the latent group variable (c) were set to the fraction of cells with mutant read counts greater than zero.

2. E-Step: Compute the posterior probability of ci = 1 for each cell, using Bayes’ rule:

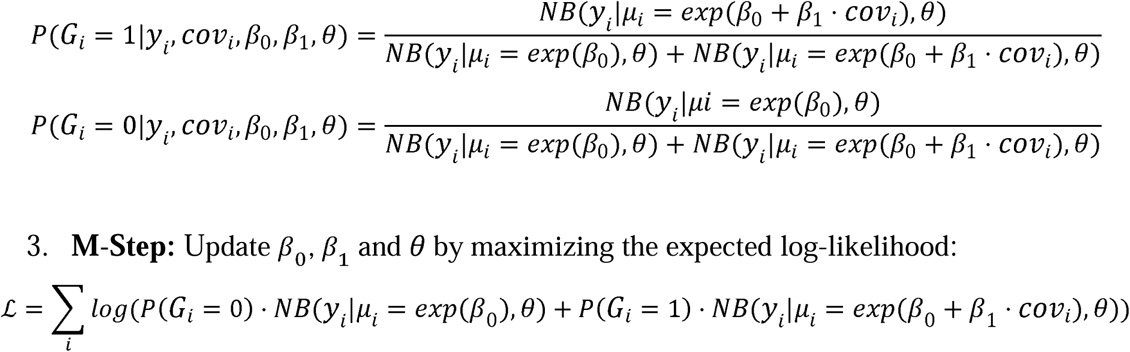

4. Iteration: Alternate between the E-step and M-step until convergence, defined as the relative change in parameter estimates falling below a predefined threshold.

The model was implemented in R. Parameter estimation was performed using the L-BFGS-B algorithm in the optim function, which supports parameter bounds. Upper bounds were set to. To discourage overestimation of the dispersion parameter, a large penalty term (106) was added to the negative log-likelihood when θ>5.

After convergence, the mutation–cell probability matrices (m by,) were merged with the CNV posterior probability matrices from Numbat. Probabilities were capped at 0.99 and floored at 0.01 before being provided as input to ScisTree, which outputs both subclonal assignments and the corresponding trees.

#### Differential expression analysis

For the RNA layer, differential gene expression analysis was performed using a pseudobulk approach with DESeq2 (*80*) to reduce false discoveries and identify common transcriptional changes across patients (*81*). For comparisons between HSC/MPP-like uMRD and donor-derived normal cells, single-cell RNA counts from CR samples were aggregated (“pseudobulked”) by patient ID, cell source (normal or uMRD), and cell type annotation. Genes with zero counts in more than five pseudobulk samples were excluded. The design gene_i_ ∼ patient ID + cell type + source was used to control for inter-patient variability (patient ID) and cell type variability, while testing for differential expression associated with the cell source (source). For comparisons between HSC/MPP-like relapse and preSCT MDS cells, ATAC and RNA counts from malignant cells were aggregated by patient ID, timepoints, and annotation. DESeq2 was used to determine the effect of disease status (preSCT or relapse) on gene expression, while EdgeR (*82*) was applied to identify differentially accessible chromatin peaks. For gene set enrichment analysis (GSEA) (*83*), gene sets were obtained using the msigdbr function in R. Analysis was performed with the fgsea package with genes ranked by fold change. For transcription factor activity differences between preSCT and relapse inferred from the ATAC layer, differential peaks from EdegeR results were subsequently analyzed for enriched transcription factor motifs using FindMotifs function in Signac (*69*).

For the protein layer, differential protein expression analysis was performed using negative binomial generalized linear models with glm.nb function in R. Antibody-derived tag (ADT) count matrices were modeled with isotype control ADT counts, patient ID, and cell source (normal or uMRD) as covariates. The effect of cell source on expression was quantified as the log_2_ fold change corresponding to the coefficient for cell source.

#### Gene regulatory network reconstruction and pathway activity analysis

Regulatory networks were inferred using SCENIC+ with default parameters, and transcriptional topics were identified with pycisTopic. Regulon activity scores were calculated using AUCell. Cytokine activity was assessed using CytoSig, with either single-cell or pseudobulked profiles as input. For cell cycle analysis, the CellCycleScoring function in the Seurat package was used to assign each cell to a cell cycle phase.

#### Bulk whole genome sequencing

*Preprocessing of BAM files*

BAM files were preprocessed following the NCI Genomic Data Commons best practices, as described at https://docs.gdc.cancer.gov/Data/Bioinformatics_Pipelines/DNA_Seq_Variant_Calling_Pipeline/. Adapter trimming was performed using Trimmomatic v0.39 with the following parameters: ILLUMINACLIP:NexteraPE-PE.fa:2:30:10 to remove Nextera paired-end adapter sequences (allowing up to 2 mismatches in the seed, a palindrome clip threshold of 30, and a simple clip threshold of 10); LEADING:3 and TRAILING:3 to trim low-quality bases from the ends of reads (Phred score <3); SLIDINGWINDOW:4:15 to remove regions where the average quality within a 4-base window dropped below 15; and MINLEN:36 to discard reads shorter than 36 bases after trimming. Trimmed reads were then aligned to the reference genome hg38 using BWA MEM v0.7.17 and converted to BAM format. Duplicate reads were removed using Picard’s MarkDuplicates v2.27.4. Finally, GATK v3.7 was used for indel realignment, utilizing known indels from dbSNP v138, the 1000 Genomes Project and Mills gold standard indels, and the GATK resource bundle, followed by base quality score recalibration to enhance BAM files for accurate variant calling.

#### Single-nucleotide variant (SNV) calling

Recipient-derived somatic SNVs were called from preprocessed BAM files using MuTect2 GATK v4.4, MuSE v2.0, and Strelka2 v2.9.10 in joint tumor-normal calling mode, with only variants identified by at least two callers being selected for downstream analyses. At pre-SCT timepoints, BAMs from bone marrow aspirates (containing exclusively recipient-derived DNA) were used as tumor samples, and BAMs from expanded T cells isolated from patient pre-SCT PBMCs were used as matched normal controls. At relapse, bone marrow aspirates contain both recipient- and donor-derived DNA. Therefore, we employed a “double-calling” strategy. First, bone marrow aspirate BAMs from relapse were called against expanded T cell BAMs from pre-SCT PBMCs (recipient germline) as above. Next, relapse BAMs were called against expanded T cell BAMs from patient postSCT PBMCs (donor germline). The intersection of the two sets of calls – one containing recipient somatic and donor somatic / germline variants, and the other containing recipient somatic / germline and donor somatic variants – effectively excludes recipient and donor germline variants, isolating recipient and donor somatic variants. We consider donor somatic variants to be biologically insignificant.

#### Identification of copy number variation

Copy number analysis was performed using the ASCAT v3.2.0 R package (*84*), with recipient T cells as matched normal controls. The ‘ascat.prepareHTS()’ function was used with default parameters to generate required LogR and B-allele frequency (BAF) files from the same BAM files as described above. After a LogR correction step, data was segmented by running ‘ascat.aspcf()’ with default parameters. Finally, tumor ploidy and purity were estimated using the ‘ascat.runAscat()’ function.

#### Variant clustering and quantification of cancer cell fraction

Variant clustering and calculation of cancer cell fraction (CCF) were performed by PyClone (*85*). PyClone requires as its input a table containing a mutation ID, the sample of origin, counts of the reference and alternate alleles, copy number information at the variant’s position, and tumor purity. Counts of the reference and alternate alleles were extracted from processed BAM files using mpileup function in samtools v1.16.1. Copy number information was obtained from ASCAT as described above (or set to default if normal karyotype). In patients and timepoints with known clonal copy number variations (CNVs) (P04, P05, P07, P08, and P09 (preSCT only)) tumor purity estimation was obtained from ASCAT output. For patients lacking clonal CNVs (i.e. subclonal CNVs or normal karyotype), tumor purity estimation was performed by calculating the VAF of the most prevalent known leukemic driver mutation and multiplying by two (assuming heterozygosity). Variants were then clustered using PyClone with default inputs except initial clusters set to 15, resulting in clusters of mutations with similar CCFs (PyClone outputs a single representative CCF for each cluster, rather than individual CCFs for each variant). Clusters with less than 10 mutations were excluded from further analyses. Visualization of changes in cancer cell fraction by mutation cluster were accomplished using the TimeScape package in R. To focus on coding somatic mutations, variant calls were annotated using ANNOVAR, then subset to exclude non-exonic variants as well as exonic synonymous variants. CCFs of exonic mutations were determined by the CCF of the PyClone cluster they were identified in prior to subsetting.

Supplementary Text Figs. S1 to S16 Tables S1 to S2 References (1–85)

