## Supplementary material for "Molecular and cellular dynamics of measurable residual disease progression in myelodysplastic syndromes": supp. figs.

### Supplementary figures

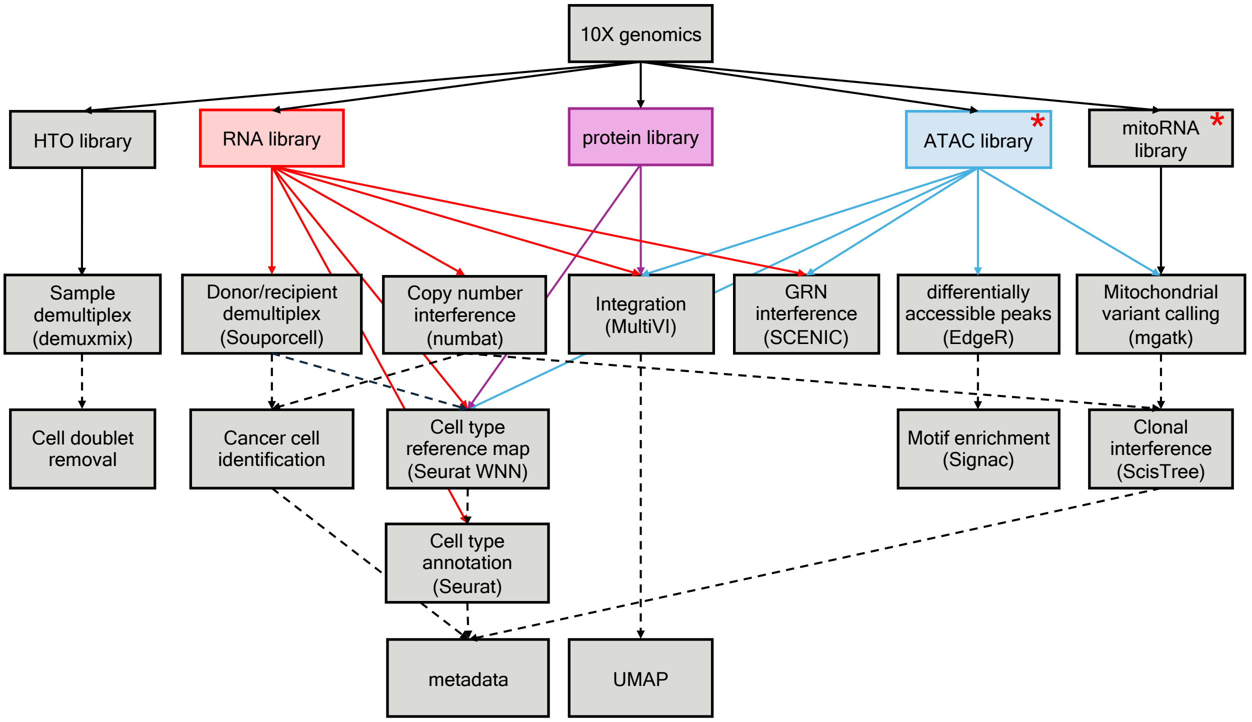


**Fig. S1. Single-cell multi-omic analysis workflow**. 10x Genomics data were processed across multiple layers, including RNA, protein, ATAC, hashtag oligo (HTO), and mitochondrial RNA libraries. *ATAC libraries were only processed from multiome kit and mitochondrial RNA libraries were only generated from 3'v3.1 kit.

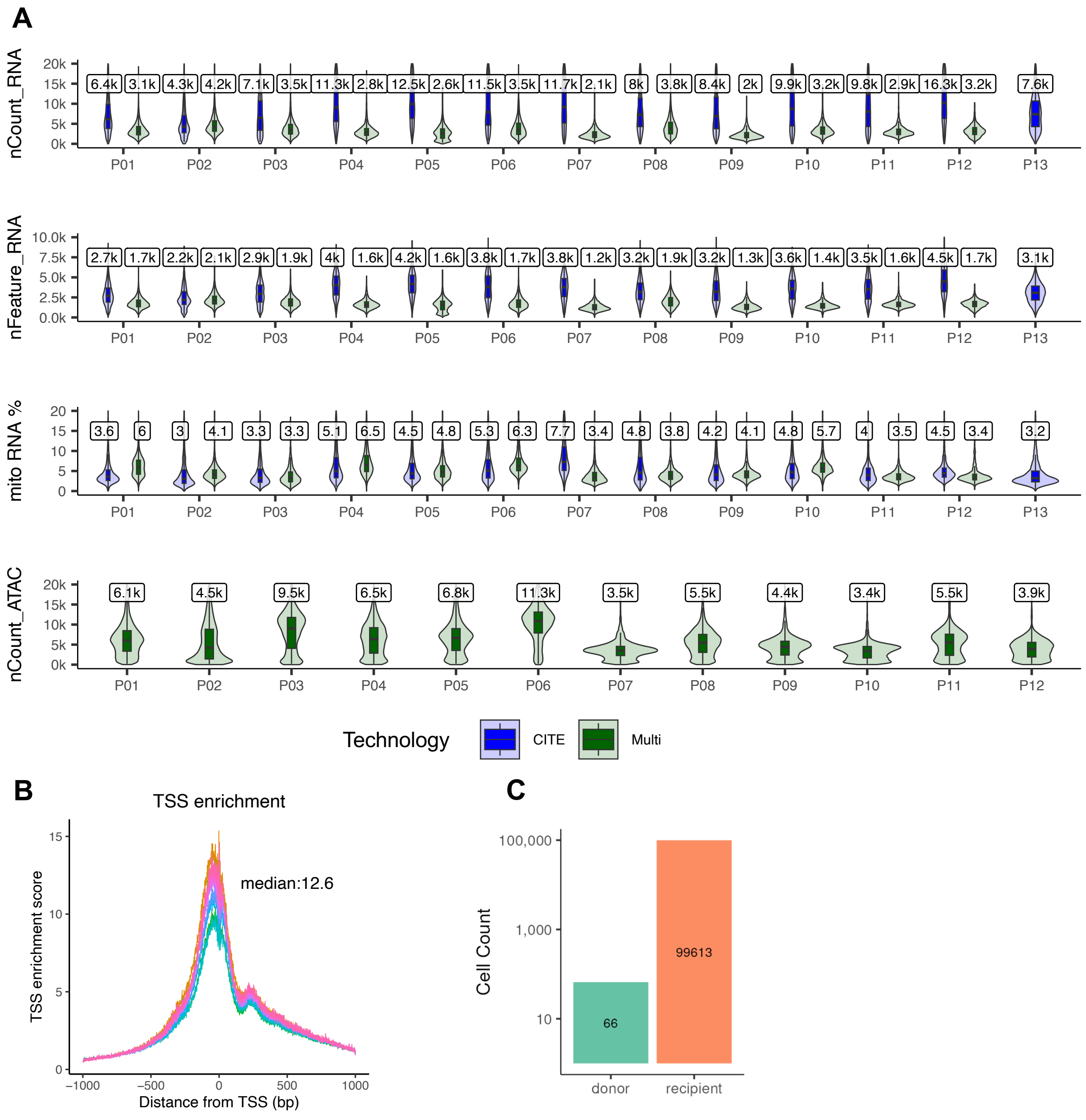


**Fig. S2. Quality control metrics of single-cell multi-omic data across patients.**
(A) Violin plots display per-cell distributions of sequencing metrics stratified by patient and technology platform. Top to bottom: total RNA UMI counts per cell (nCount_RNA), number of detected RNA features (nFeature_RNA), percentage of mitochondrial RNA transcripts (mito_RNA%), and total ATAC fragment counts per cell (nCount_ATAC). Each panel is colored by library technology: CITE-seq (blue) and MultiVI (green). Median values are indicated above each distribution. (B) Transcription start site (TSS) enrichment plot showing aggregate TSS accessibility profiles across all cells. The median TSS enrichment score is 12.6. (C) Barplot shows the number of cells assigned as donor (n = 66) or recipient (n = 99,613) from pre-transplant (preSCT) samples, in which only recipient cells are expected. The presence of donor-assigned cells reflects misclassification, yielding an estimated Souporcell error rate of 0.06% calculated as 66 / (66 + 99,613)

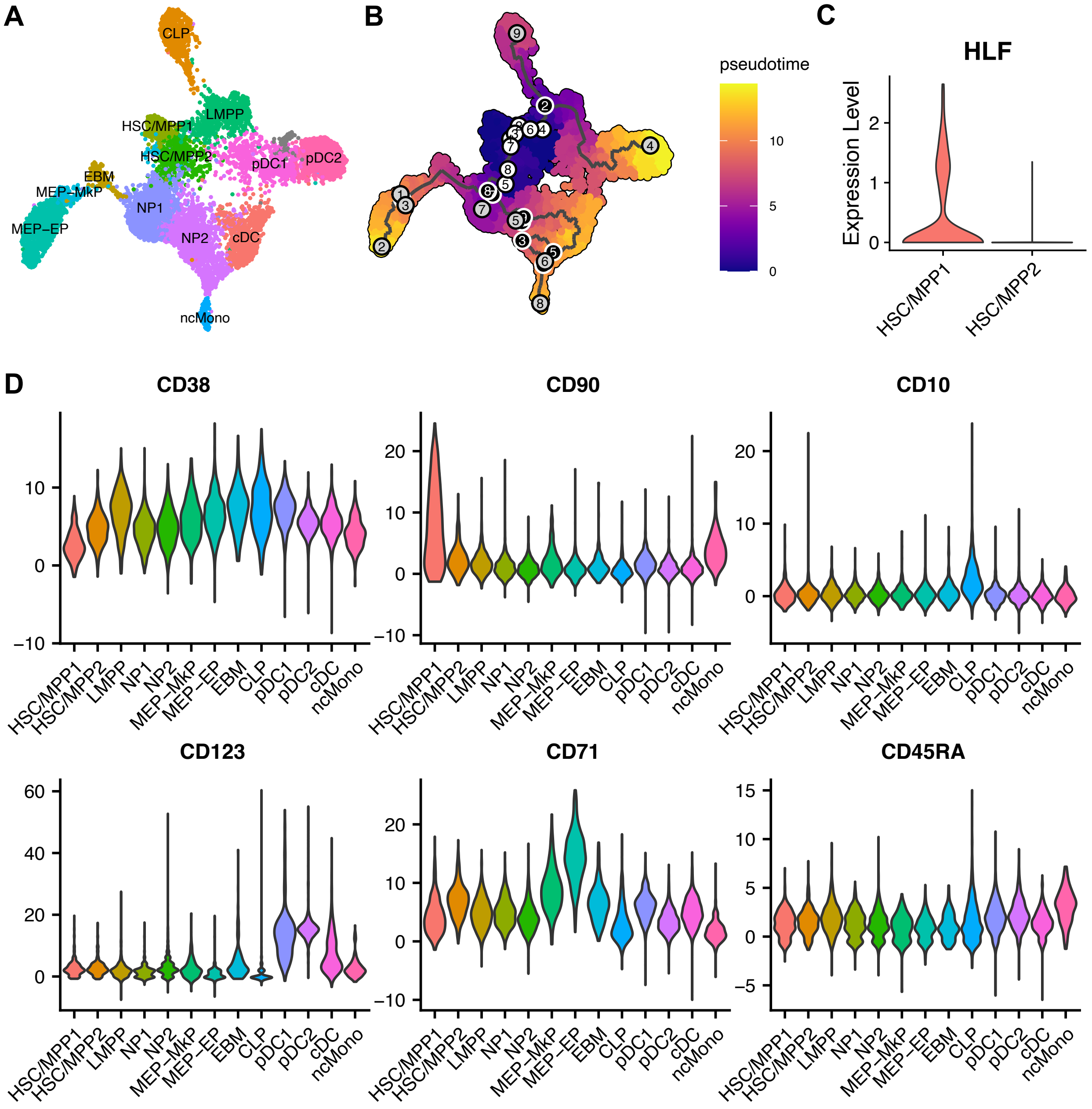


**Fig. S3. Reference map of hematopoietic stem and progenitor cell (HSPC) states**.
(A) UMAP projection of reference CD34⁺ cells showing trimodally defined HSPC subpopulations. (B) Diffusion pseudotime trajectory overlaid on the reference map, illustrating inferred differentiation trajectories using Monocle 3. (C) HLF RNA expression in HSC/MPPs. (D) Violin plots showing normalized surface protein expression of canonical markers (CD38, CD90, CD10, CD123, CD71, CD45RA) across annotated reference cell types. **HSC**: Hematopoietic stem cells, **MPP**: Multipotent progenitor cells, **NP**: Neutrophil progenitors, **pDC**: Plasmacytoid dendritic progenitors, **ncMono**: non-classic monocytes, **MkP**: Megakaryocyte progenitor, **EP**: Erythroid progenitors, **EBM**: Eosinophil-basophil-mast cell progenitors, **CLP**: Common lymphoid progenitors.

**A**

| | HSC/MPP1 | HSC/MPP2 | LMPP | NP1 | NP2 | cDC | ncMono | CLP | pDC1 | pDC2 | EBM | MEP-EP | MEP-MkP |
|---|---|---|---|---|---|---|---|---|---|---|---|---|---|
| HSCs & MPPs | 80 | 679 | 21 | 0 | 0 | 0 | 0 | 1 | 0 | 0 | 0 | 1 | 23 |
| Lymphomyeloid prog | 0 | 409 | 16 | 0 | 0 | 0 | 0 | 19 | 1 | 0 | 0 | 3 | 4 |
| Early promyelocytes | 0 | 279 | 2 | 84 | 0 | 1 | 0 | 2 | 1 | 0 | 0 | 0 | 0 |
| Late promyelocytes | 0 | 111 | 0 | 67 | 22 | 175 | 1 | 0 | 2 | 0 | 0 | 2 | 0 |
| Conventional dendritic cell 1 | 0 | 25 | 2 | 0 | 0 | 187 | 0 | 0 | 19 | 2 | 0 | 2 | 0 |
| Megakaryocyte progenitors | 0 | 28 | 0 | 0 | 0 | 0 | 0 | 0 | 0 | 0 | 0 | 55 | 98 |
| Non-classical monocytes | 0 | 0 | 0 | 0 | 0 | 1 | 96 | 0 | 0 | 0 | 0 | 0 | 0 |
| Pre-B cells | 0 | 7 | 0 | 0 | 0 | 0 | 0 | 181 | 0 | 0 | 0 | 1 | 0 |
| Pro-B cells | 0 | 17 | 4 | 0 | 0 | 0 | 0 | 494 | 0 | 0 | 0 | 0 | 0 |
| Pre-pro-B cells | 0 | 1 | 4 | 0 | 0 | 0 | 0 | 17 | 0 | 0 | 0 | 0 | 0 |
| Small pre-B cell | 0 | 0 | 0 | 0 | 0 | 0 | 0 | 12 | 0 | 0 | 0 | 0 | 0 |
| Plasmacytoid dendritic cell progenitors | 0 | 20 | 20 | 0 | 0 | 3 | 0 | 5 | 92 | 5 | 0 | 0 | 0 |
| Plasmacytoid dendritic cells | 0 | 0 | 1 | 0 | 0 | 1 | 0 | 0 | 3 | 98 | 0 | 0 | 0 |
| Eosinophil-basophil-mast cell progenitors | 0 | 8 | 1 | 0 | 0 | 0 | 0 | 0 | 0 | 0 | 44 | 5 | 1 |
| Erythro-myeloid progenitors | 0 | 46 | 1 | 0 | 0 | 0 | 0 | 0 | 0 | 0 | 0 | 36 | 131 |
| Early erythroid progenitor | 0 | 13 | 0 | 0 | 0 | 0 | 0 | 0 | 0 | 0 | 0 | 320 | 0 |
| Late erythroid progenitor | 0 | 2 | 0 | 0 | 0 | 0 | 0 | 0 | 0 | 0 | 0 | 316 | 0 |
| Megakaryocyte progenitors | 0 | 28 | 0 | 0 | 0 | 0 | 0 | 0 | 0 | 0 | 0 | 55 | 98 |

**B**

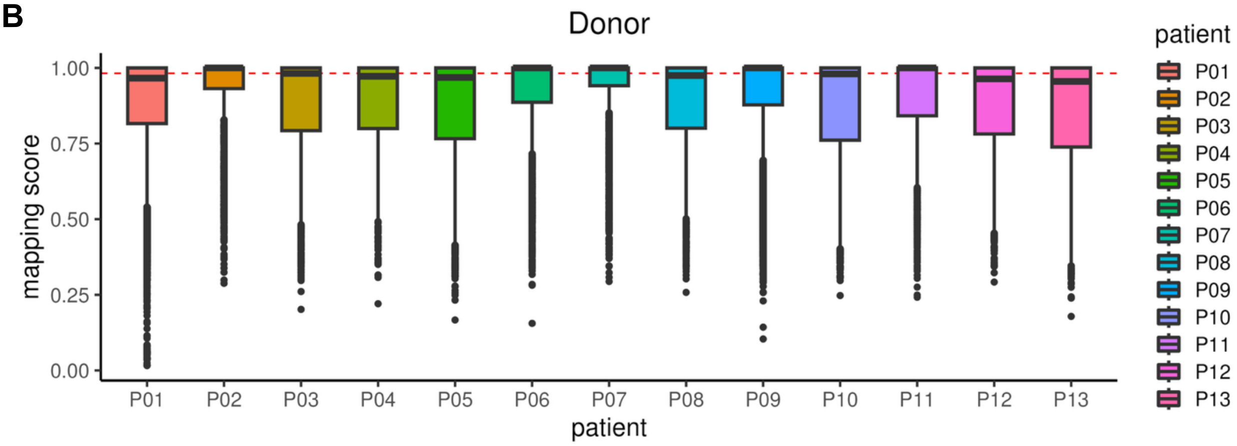


**C**

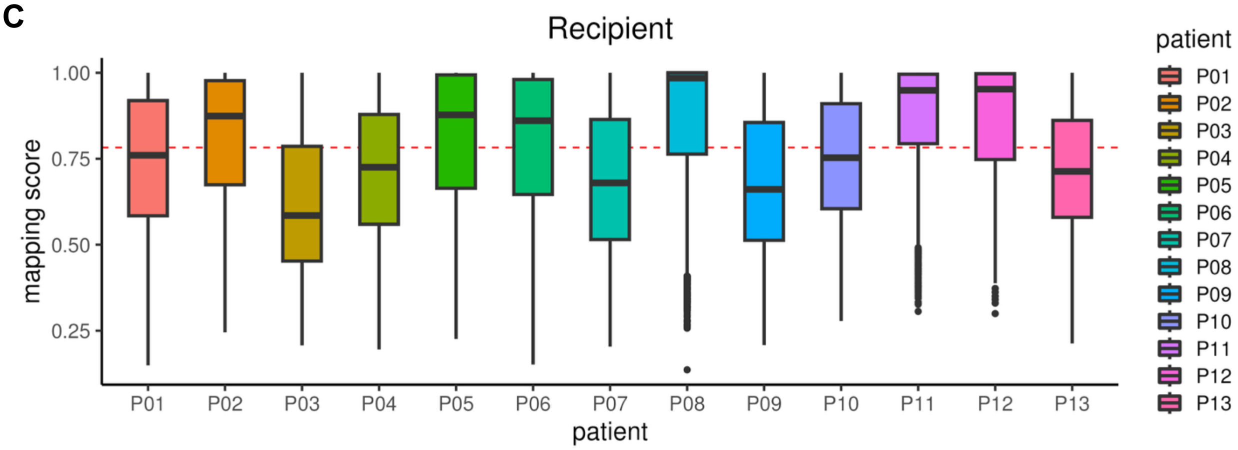


**Fig. S4**. **Quality control of reference HSPC map and mapping score of donor- and recipient-derived cells to the map.** (A) Cross-validation of cell type annotation and label transfer by mapping cells from the reference atlas (S. Triana *et al*., Nature Immunology 2021 (*1*); blue, left) onto our reference map (green, top). The resulting correspondence matrix shows high consistency between the two annotation schemes, confirming the robustness of our label transfer. (B-C) Boxplots showing the mapping scores of CD34$^{+}$ cells from each patient for (B) donor-derived and (C) recipient-derived compartments. Mapping scores reflect the consistency of individual cells to reference cell types, with higher scores indicating more confident annotation. Each box represents one patient, colored accordingly. The red dashed line indicates the median mapping score across all donor- or recipient-derived cells, respectively.

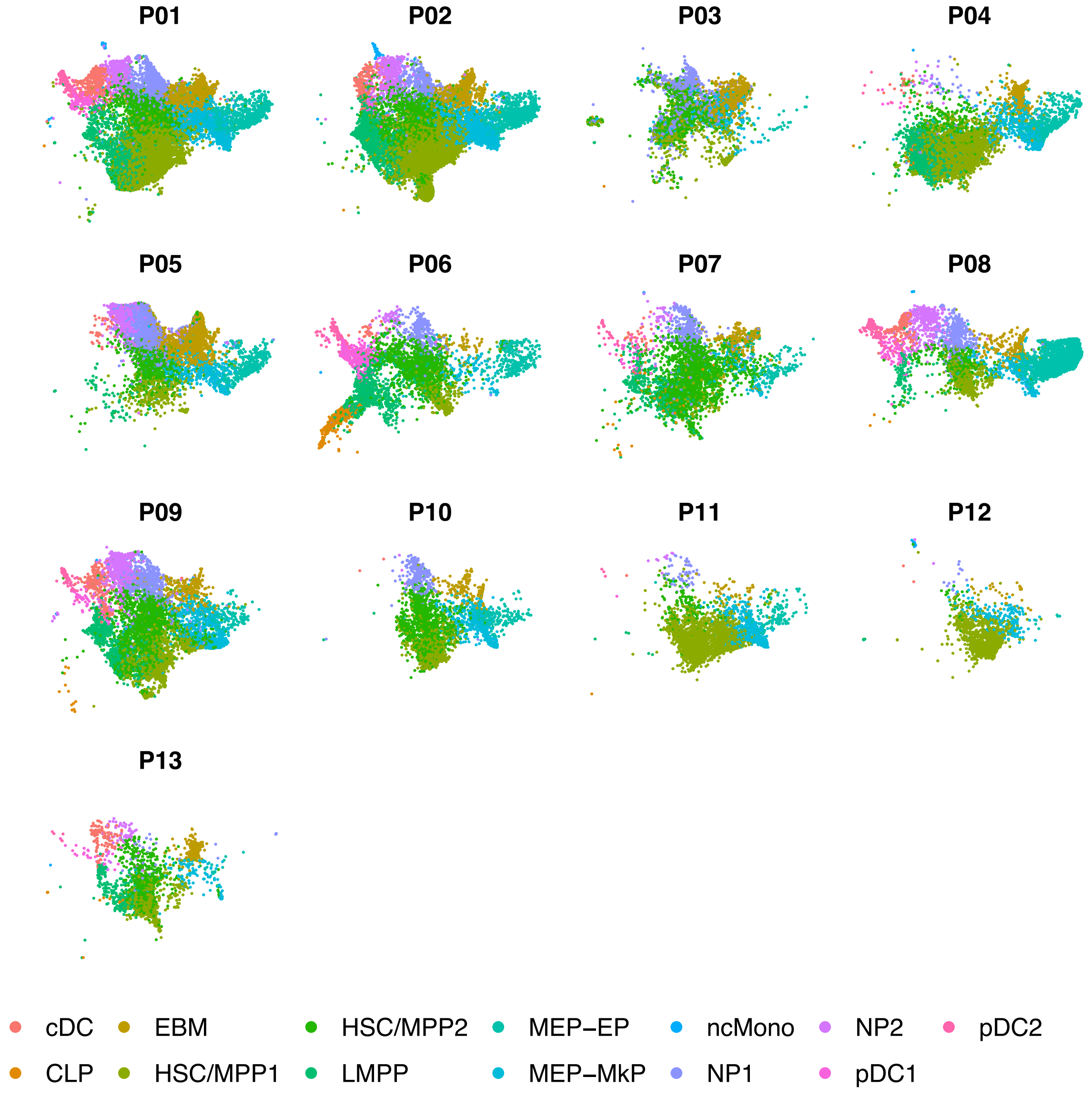


**Fig. S5**. **UMAP projections of recipient-derived CD34$^+$ cells from 13 patients (P01–P13)**. Each point represents a single cell, colored by cell type annotation: **HSC**: Hematopoietic stem cells, **MPP**: Multipotent progenitor cells, **NP**: Neutrophil progenitors, **pDC**: Plasmacytoid dendritic progenitors, **ncMono**: non-classic monocytes, **MkP**: Megakaryocyte progenitor, **EP**: Erythroid progenitors, **EBM**: Eosinophil-basophil-mast cell progenitors, **CLP**: Common lymphoid progenitors.

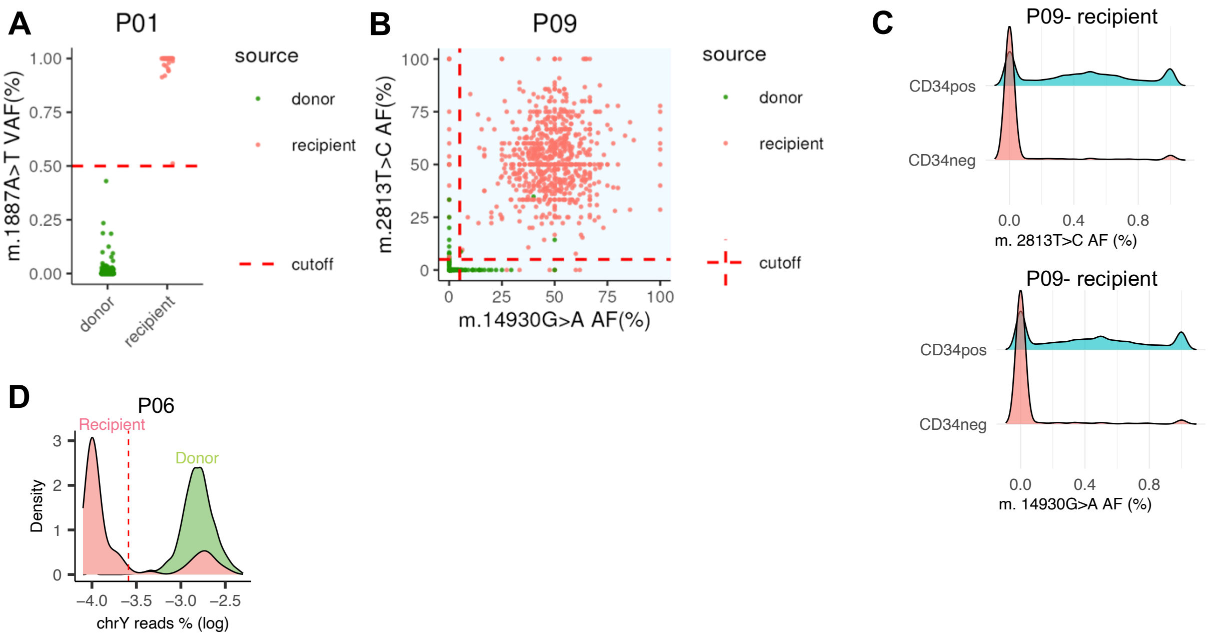


**Fig. S6. Identification of malignant recipient-derived cells using mitochondrial variants and loss of chromosome Y.** (A) Variant allele frequency (VAF) of mitochondrial variant m.8393T>C in donor (green) and recipient (red) cells from patient P01. (B) Scatter plot of VAFs for two mitochondrial variants (m.14930G>A vs. m.8393T>C) in patient P09, with each point representing a single cell, color-coded by inferred genotype. (C) Distribution of the two mitochondrial variants in $CD34^+$ and $CD34^-$ compartments of patient P09, showing enrichment of the variants in the $CD34^+$ population. (D) Density distribution of chromosome Y (chrY) read percentage in donor (green) and recipient (red) cells. Red dashed line indicates the classification threshold separating normal from malignant cells. The 99% expression or VAF interval from donor reference cells was used as a cutoff, thereby controlling the false positive rate for abnormal cell detection at approximately 1%, under the assumption that donor cells represent normal expression variability.

A

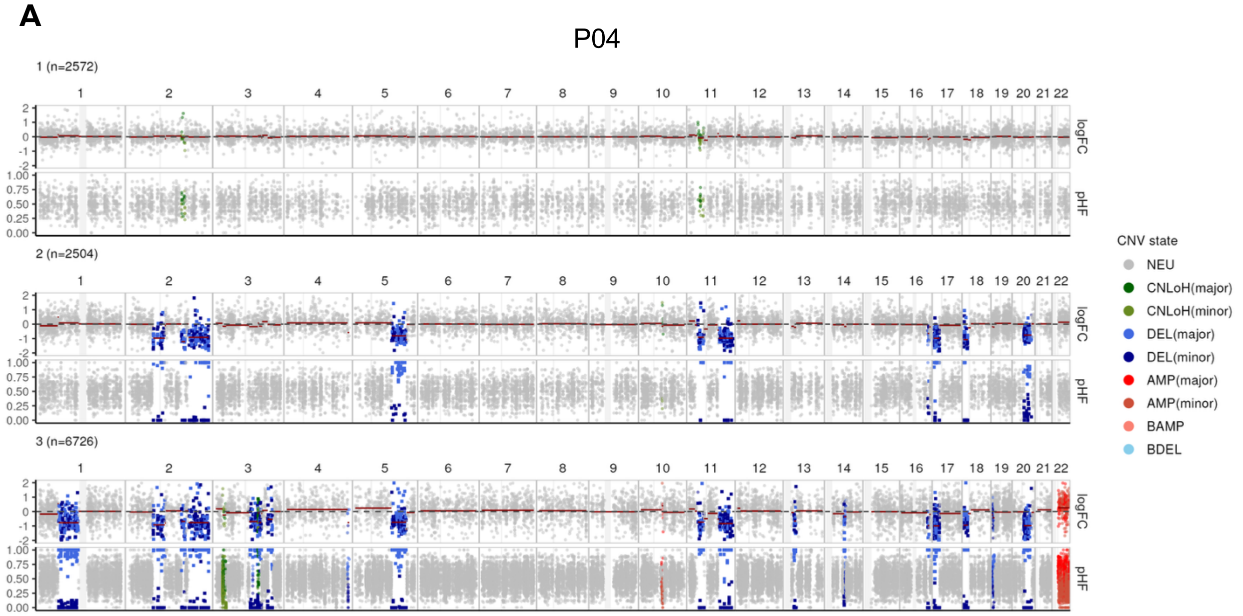
P04
1 (n=2572)
2 (n=2504)
3 (n=6726)
1 2 3 4 5 6 7 8 9 10 11 12 13 14 15 16 17 18 19 20 21 22
logFC
pHF
CNV state
NEU
CNLoH(major)
CNLoH(minor)
DEL(major)
DEL(minor)
AMP(major)
AMP(minor)
BAMP
BDEL

B

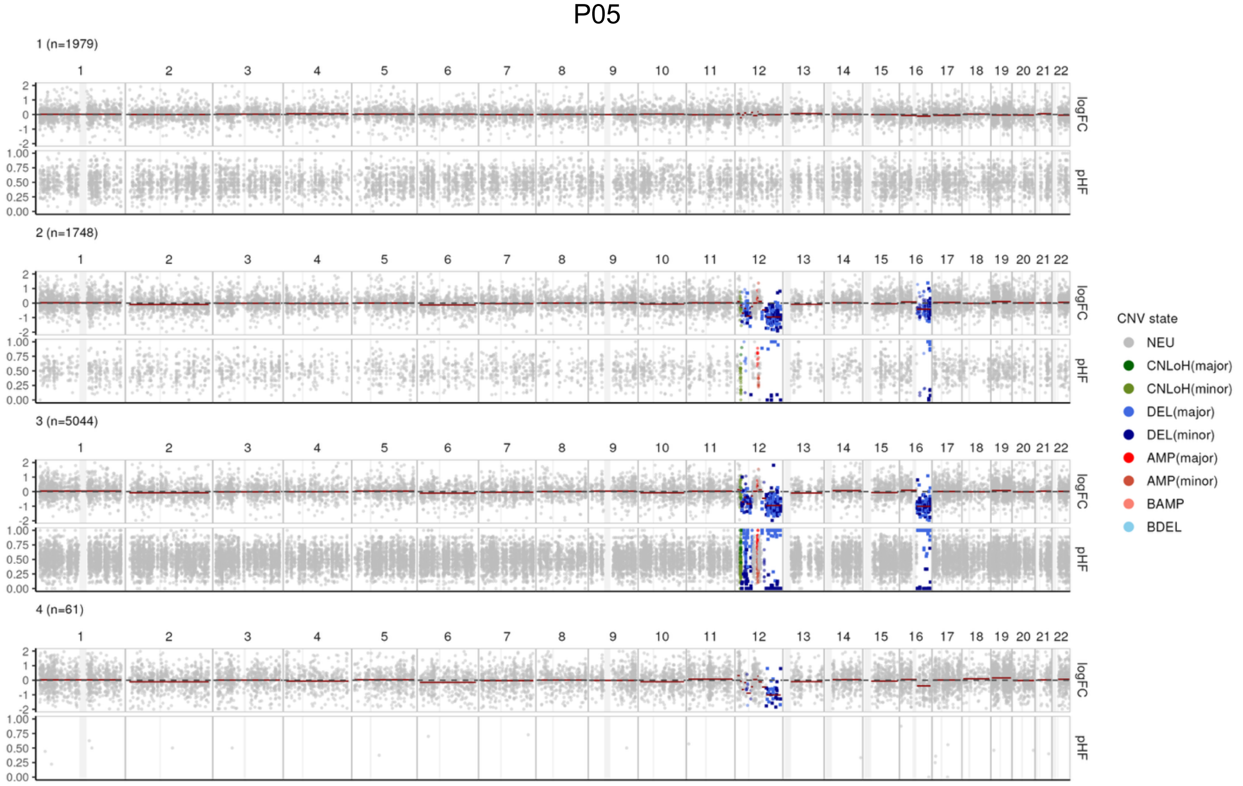
P05
1 (n=1979)
2 (n=1748)
3 (n=5044)
4 (n=61)
1 2 3 4 5 6 7 8 9 10 11 12 13 14 15 16 17 18 19 20 21 22
logFC
pHF
CNV state
NEU
CNLoH(major)
CNLoH(minor)
DEL(major)
DEL(minor)
AMP(major)
AMP(minor)
BAMP
BDEL

**C**

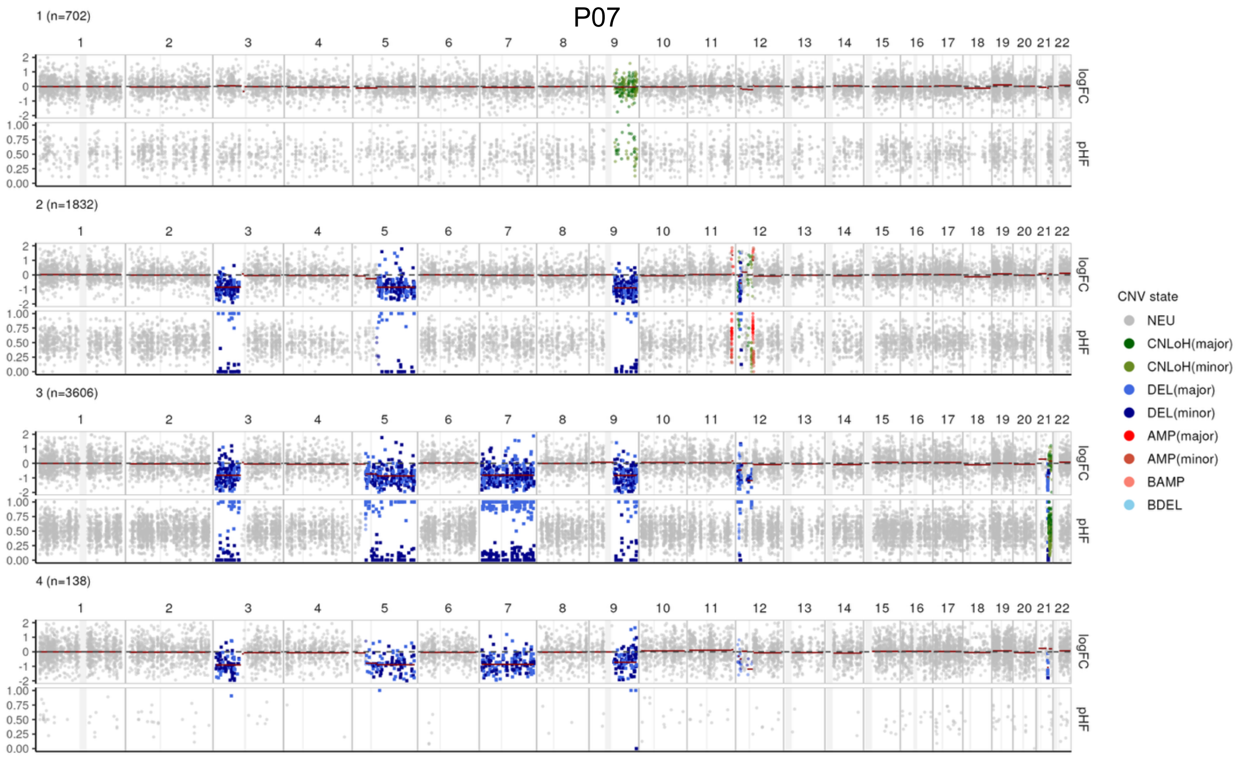


**D**

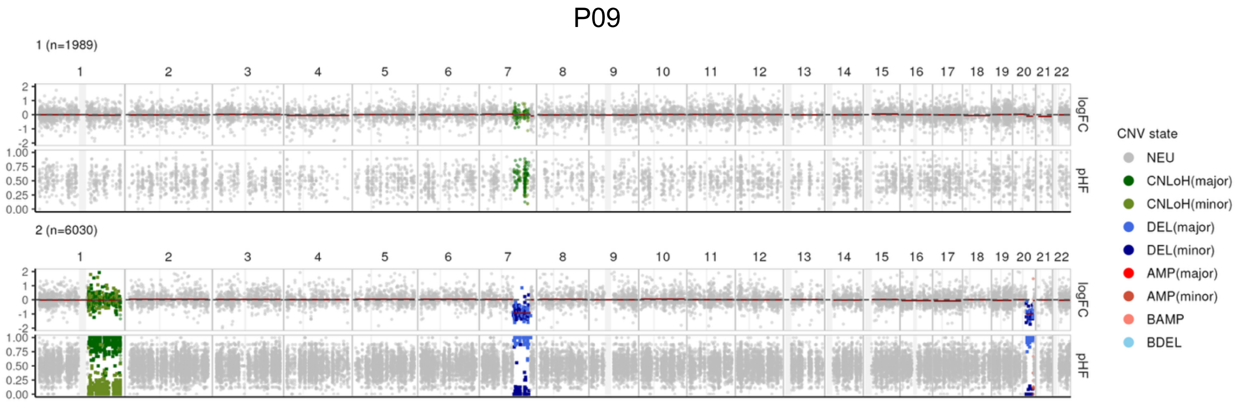


**Fig. S7. Inference of copy number alterations (CNAs) in single cells using Numbat for identification of malignant recipient-derived cells across patients.** (A–D) Pseudobulk CNA profiles for patient P04 (A), P05 (B), P07 (C), and P08 (D), showing log expression fold-change (logFC, top) and paternal haplotype frequency (pHF, bottom) across somatic chromosomes. Gray vertical bars represent centromeres and gap regions. CNV clones were identified based on shared CNA patterns; clone 1 represents cells without detectable CNAs. Copy number states are color-coded as shown.

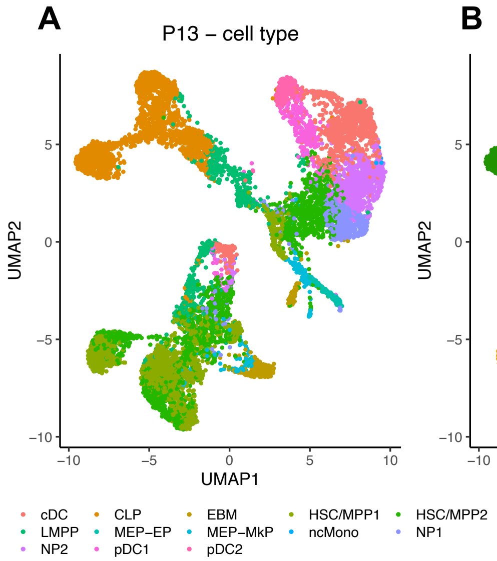


**Fig. S8. UMAP visualization and clustering of single-cell transcriptomes from patient P13.** (A) UMAP colored by cell type annotation. (B) UMAP colored by donor (green) and recipient (orange) origin inferred by Souporcell. (C) UMAP colored by clusters labeled as cancer (pink) or normal (blue). Clustering was performed at low resolution to identify broad transcriptional states. Clusters enriched for donor cells were labeled as "normal" and used to distinguish malignant from normal recipient-derived cells.

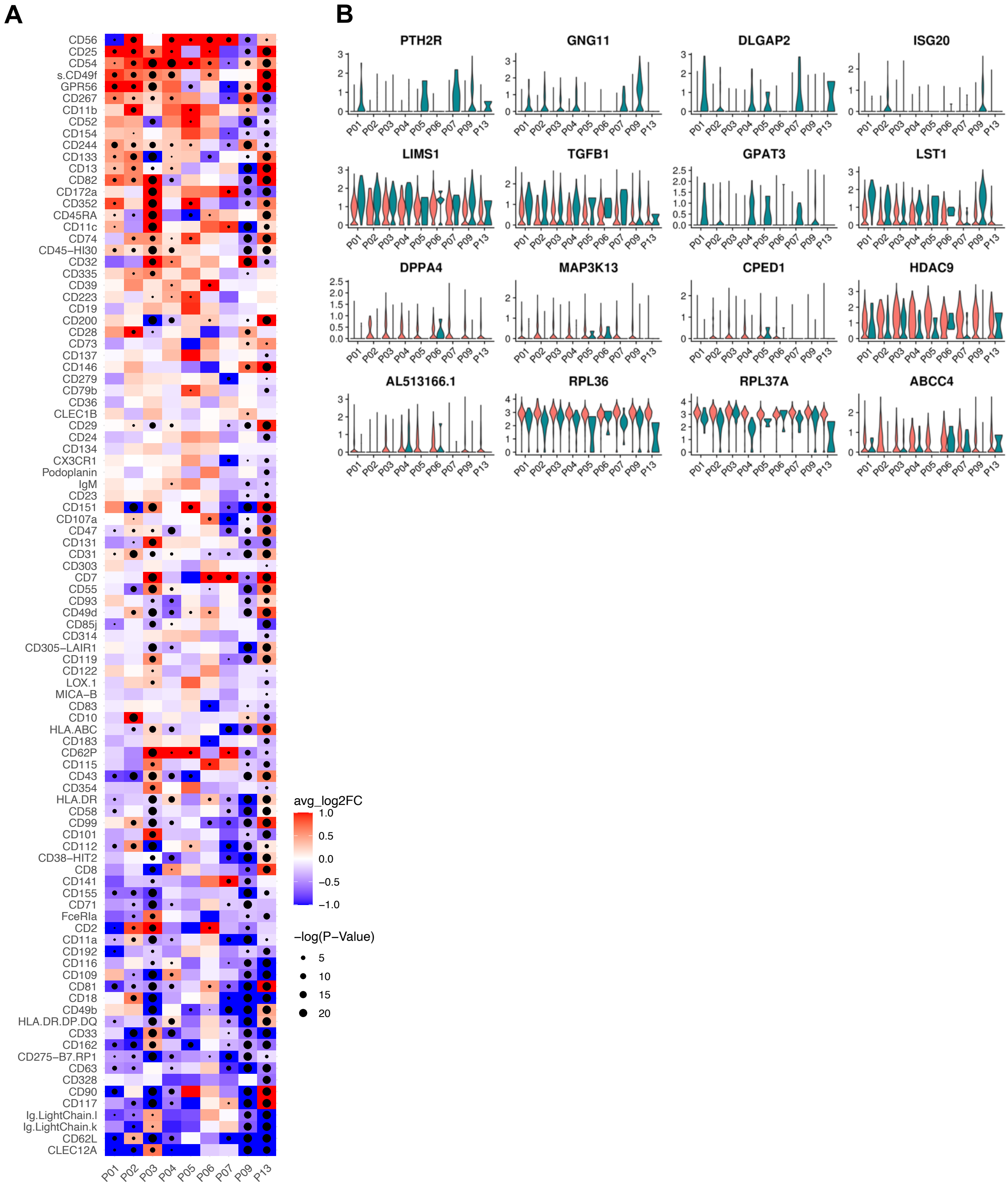


**Fig. S9. Differential surface protein and RNA expression between uMRD and normal HSC/MPP cells**. (A) Heatmap showing log fold-change in surface protein expression between uMRD and donor-derived normal cells. Red indicates upregulation in uMRD cells; blue indicates downregulation. Rows represent individual surface proteins; columns represent matched patient–cell type pairs. Statistical significance was assessed using the Wilcoxon rank-sum test; proteins with significant differences are marked with dots. (B) Violin plots showing normalized expression levels of selected genes across patients, split by donor (red) or recipient (green) cells.

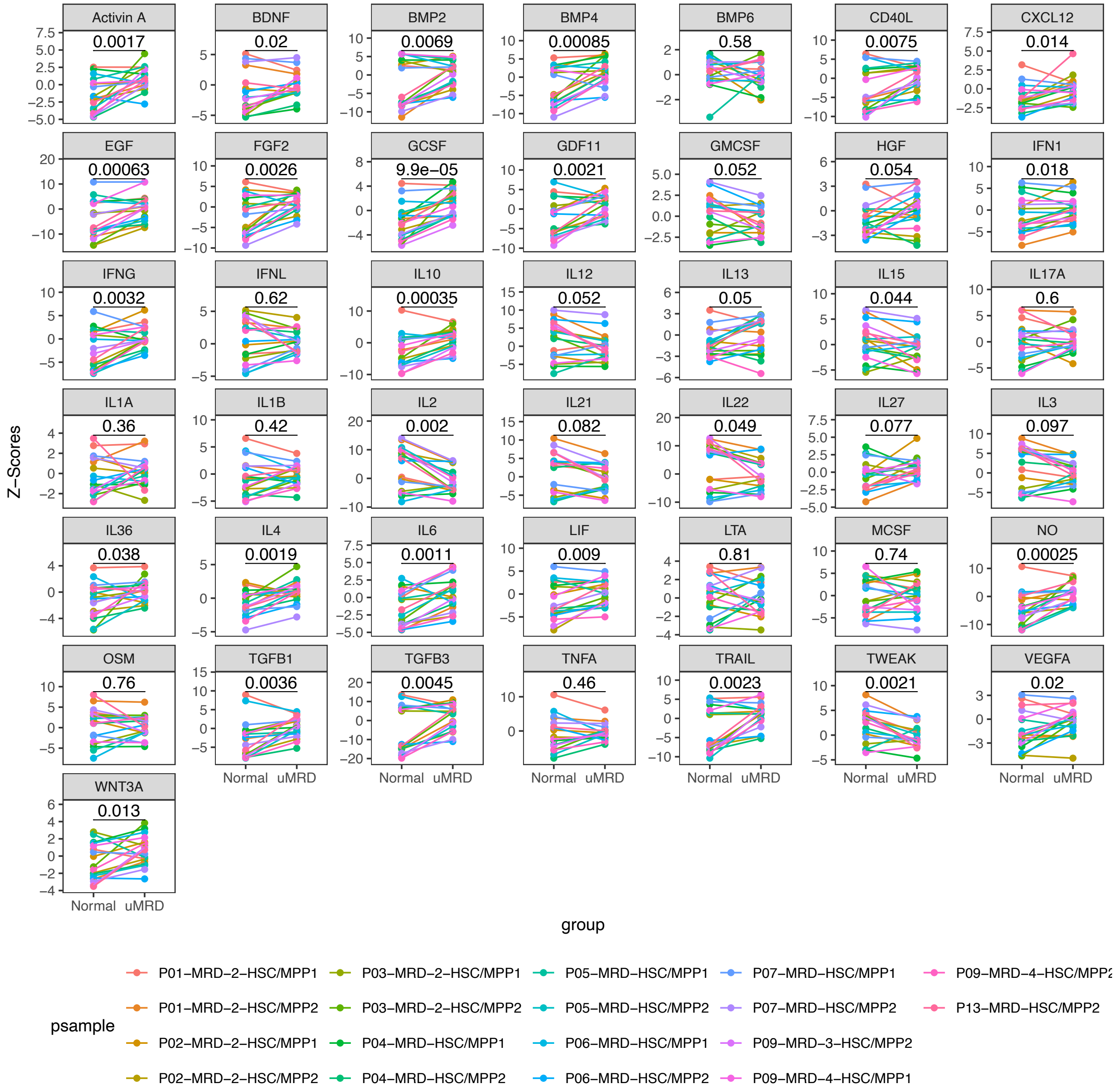


**Fig. S10. Differential cytokine and growth factor signaling between donor-derived (normal) HSC/MPP cells and HSC/MPP-like uMRD**. Signaling activity (Z-scores) was inferred using CytoSig from pseudobulk gene expression profiles, aggregated by matched patient–cell type pairs labeled as normal (donor-derived) or malignant (recipient-derived). Each line represents a patient–cell type pair (e.g., P01–HSC/MPP1, P02–HSC/MPP2). Statistical significance was assessed using paired Wilcoxon signed-rank tests. Altered signaling of factors in malignant cells highlights intrinsic dysregulation independent of bone marrow environmental differences.

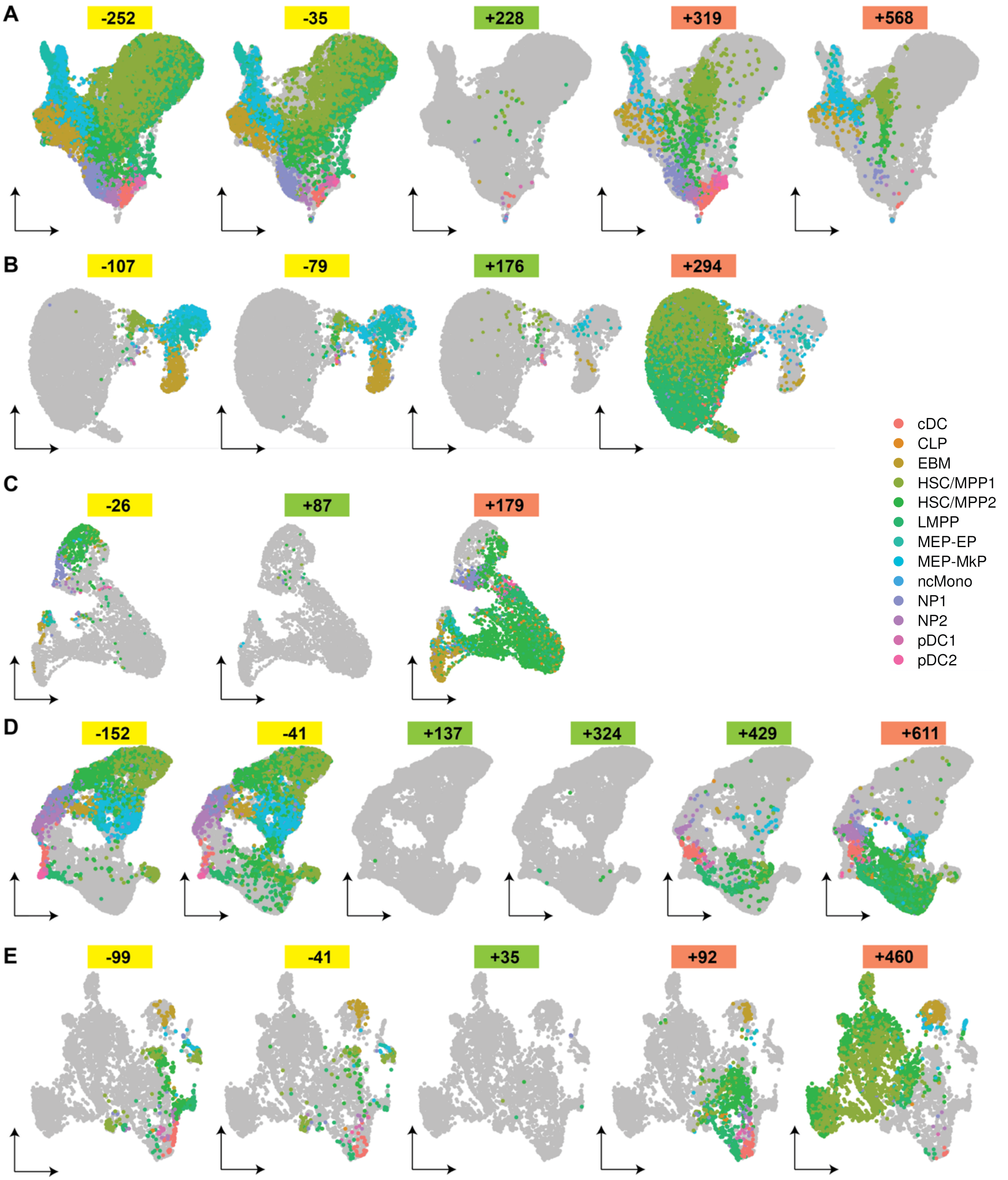


**Fig. S11. Patient-specific longitudinal MultiVI UMAPs of $CD34^+$ cells**. MultiVI-based UMAP projections of $CD34^+$ cells MDS patients P01 (A), P04 (B), P07 (C), P09 (D), and P13 (E) sampled longitudinally across key clinical timepoints: preSCT (yellow), CR (green), and relapse (red). Each panel corresponds to a single timepoint, labeled with the number of days relative to SCT (negative values = preSCT; positive values = postSCT). Cells are colored by inferred cell type.

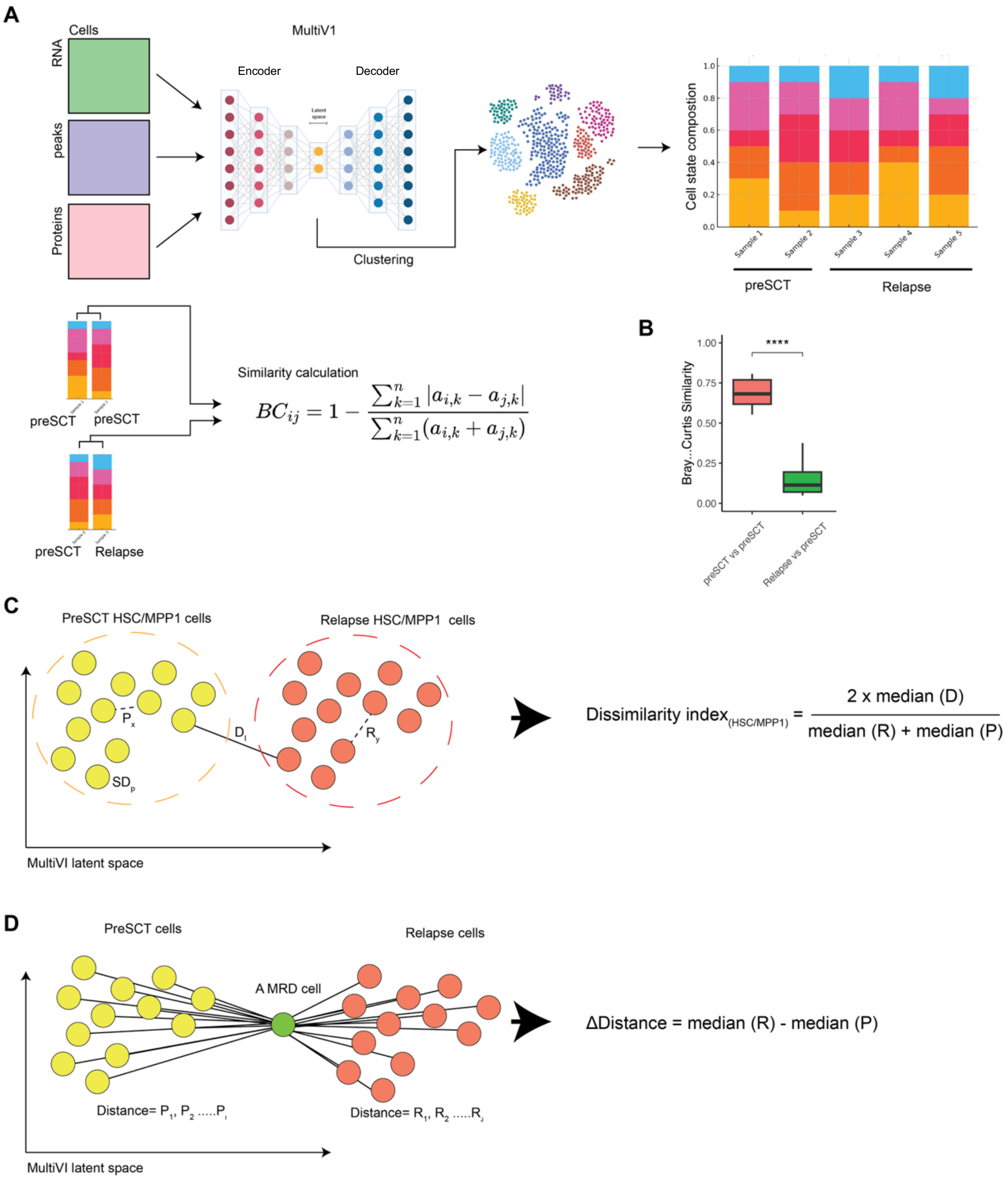


**Fig. S12. Quantification of MRD cell similarity and cell state evolution in MultiVI latent space.** (A) Overview of MultiVI-based integration of scRNA, surface protein, and cell identity data. Latent embeddings are derived via encoder-decoder architecture and subjected to clustering. The resulting clusters are defined as cell states. Stacked bar plots display cell-type composition in each sample. (B) Bray–Curtis similarity scores comparing cell state compositions across preSCT–preSCT and relapse–preSCT sample pairs. Significantly reduced similarity was observed between relapse and preSCT samples (****$P < 0.0001$, two-sided Wilcoxon test). (C) Schematic illustrating the dissimilarity index of given cell type, calculated from pairwise distances within preSCT *P*, within relapse *R*, and between preSCT–relapse *D* in latent space. (D) Visualization of ΔDistance metric: for each MRD cell, distance to preSCT cells ($P_1 \ldots P_i$) and relapse cells ($R_1 \ldots R_j$) is computed; ΔDistance is defined as the difference between median (R) and median(P), reflecting shift in phenotypic identity.

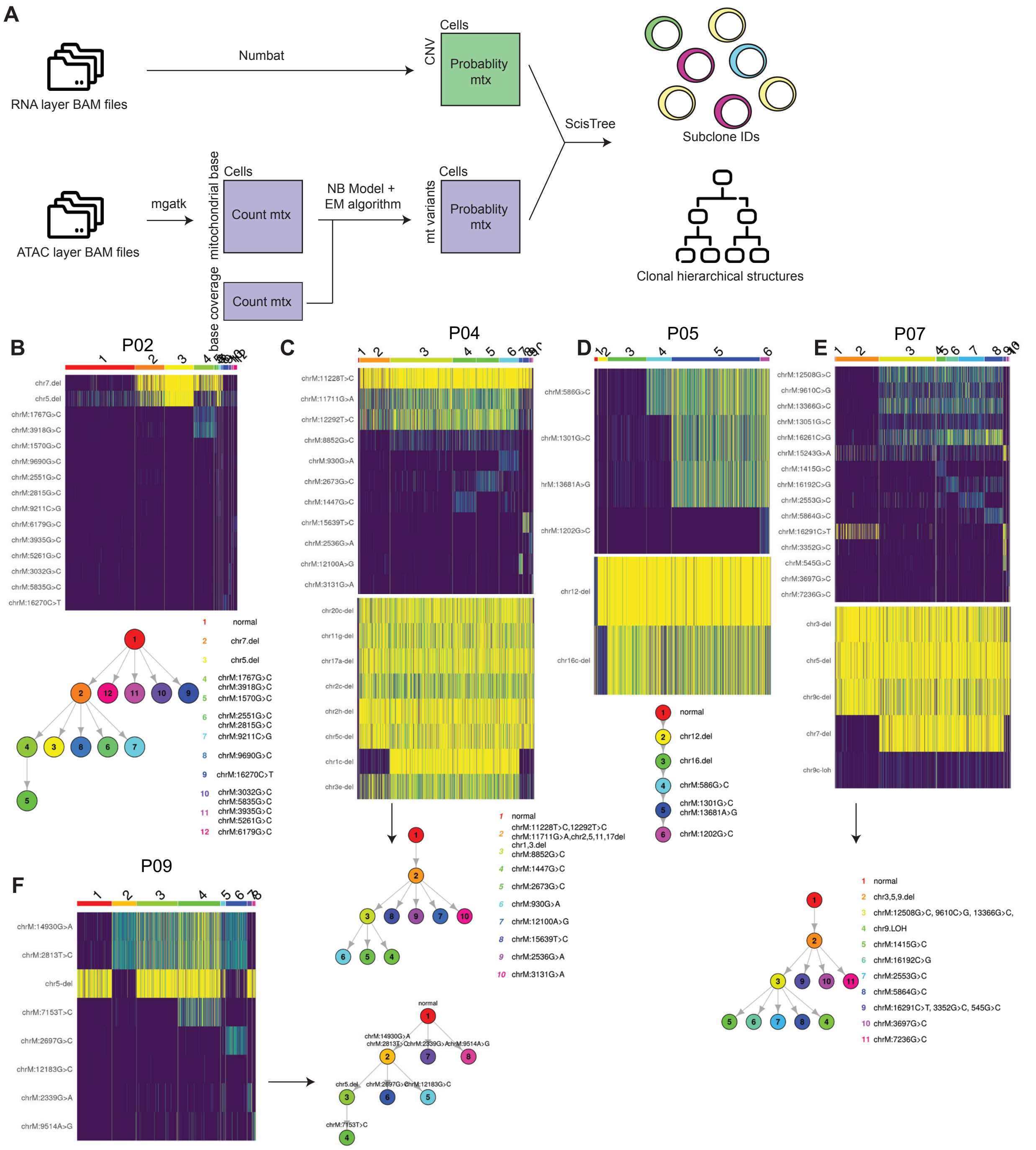


**Fig. S13 Subclonal reconstruction.** (A) Schematic of the workflow integrating RNA and ATAC layers for clonal inference. Mitochondrial genotype probabilities were estimated using a negative binomial model, and CNV posteriors were inferred with Numbat. Probability matrices were merged and provided to ScisTree (*2*) for clonal assignment and phylogenetic tree reconstruction. (B-F) Heatmaps showing mitochondrial variants and CNV states across representative patients, with corresponding phylogenetic trees depicting subclonal relationships. Colors indicate distinct subclones.

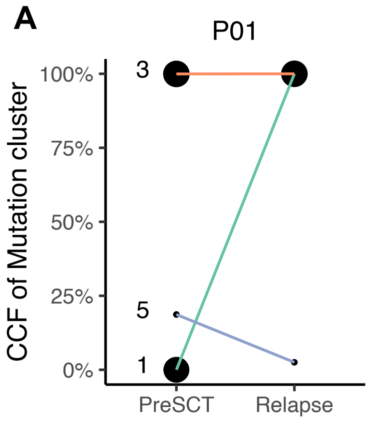


| | |
|---|---|
| 1 | KDSR, PLSCR4, PAX4 |
| 3 | MUC15, OR4A15, SYTL2 PCDH8, DHRS1, ATAD3C CFH, PARP1, TINAGL1 ASXL1, MYH9, TST, SF3B1 |
| 5 | KCNMA1, POLR1G, PPP1R12C |

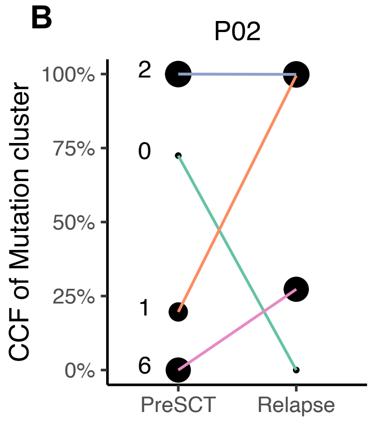


| | |
|---|---|
| 0 | SETBP1 |
| 1 | PTPN11, SYNE3, EPM2AIP1, KCNQ3 |
| 2 | HOGA1, METTL22, SUMO2, SLC14A2, U2AF2, ASXL1, SPHKAP, RFTN1, PLXNB1, POPDC3 |
| 6 | PAPSS2, TGM5, MAST1, MYO1F, WDR26, KIF16B, STAG2, GPT, KRTAP24-1 |

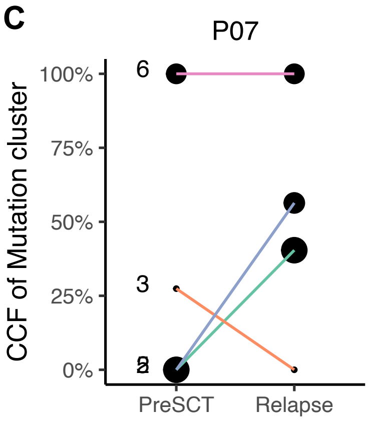


| | |
|---|---|
| 2 | ANK3, CD6, MAP6, PTPN11, GALNT9, FREM2, CHD2, SCNN1B, MVD, KRT13, SLC38A10, RYR1, OBSCN, ACTN2, GAB4, CECR2, HECW2, GLB1L, SLC16A14, USP40, EML6, TSGA10, CLRN2, MTHFD2L, PCDHA6, GRM6, GPRC6A, CUL9, COL14A1, MROH1, SCARA3, RTL8B, FAM47C, TIMM17B |
| 3 | ATP2A1, NRIP1, U2AF1, U2AF1L5, USP49, PCSK5 |
| 5 | OR5B3, CIT, AHNAK2, ZNF888, NRAS, UQCRHL, TRAF3IP3, COL6A2, MYO18B, NCK2, NXPE3, ALG1L, XPC, LAMB2, MYOZ2, OTOP1, ANKS1A |
| 6 | GRID1, SLC22A9, TSGA10IP, LRRIQ1, MYH3, ARHGAP44, TP53, GUCY2D, PPOX, LYST, PRR18, BHLHA15 |

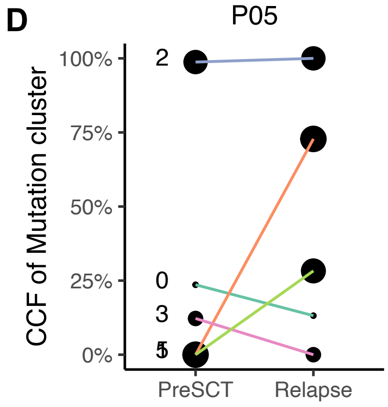


| | |
|---|---|
| 0 | ATAD3B |
| 1 | PPM1H, SIX1, BICDL2, CHD9, CHERP, ETV3, KLF18, CDCP2, RIN2, IFNAR2, TLK1, PCARE, RMDN2, TMEM45A, OR5K4, ARHGEF6 |
| 2 | ZNF248, FAM160A2, KDM2B, ANKFN1, TP53, PTGS2, CHD5, PPIL2, NEK10, UGT8, FBXO38, EN2, TTC39B |
| 3 | VWCE, TECPR1, PXDNL |
| 5 | ARID5B, SMYD4, RNF213, ATCAY, ZDHHC8, EIF5B, RELL2, ATXN1 |

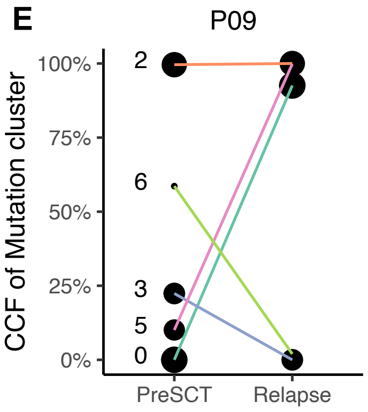


| | |
|---|---|
| 0 | TRUB1, MAT1A, LTBP3, PDE2A, OR4M1, YLPM1, LMAN1L, CACNA1H, PRRT2, FAAP100, BORCS6, PIK3R5, IRGC, TDRD5, NPHP1, SLC35F5, NEB, ASTL, PRDM9, TREML4, CCND3, TTYH3, ZMIZ2, CA3 |
| 2 | DCPS, PPM1A, UACA, ADAMTS18, SMAD7, NLRP4, CD101, NCF2, ASXL1, GATA2, ZNF654, KIAA1109, GCNT2, HIVEP2, BPHL, KMT2C |
| 3 | C11orf71 |
| 5 | TUBB8B, PLAGL1 |
| 6 | ZNF800 |

**Fig. S14 Dynamics of WGS-derived mutation clusters from preSCT to relapse**. Cellular prevalence of mutation clusters inferred from whole-genome sequencing (WGS) of sorted MDS samples at pre-transplant (PreSCT) and relapse timepoints for patient P01 (A), P02 (B), P07 (C), P05 (D), and P09 (E). Each panel shows the cancer cell fraction (CCF) of individual mutation clusters across timepoints. Clusters were derived based on mutation co-occurrence. Only clusters containing at least one exonic mutation are shown. Numbers denote cluster IDs. Accompanying tables list genes with exonic mutations within each cluster. Known AML-associated genes are highlighted in red.

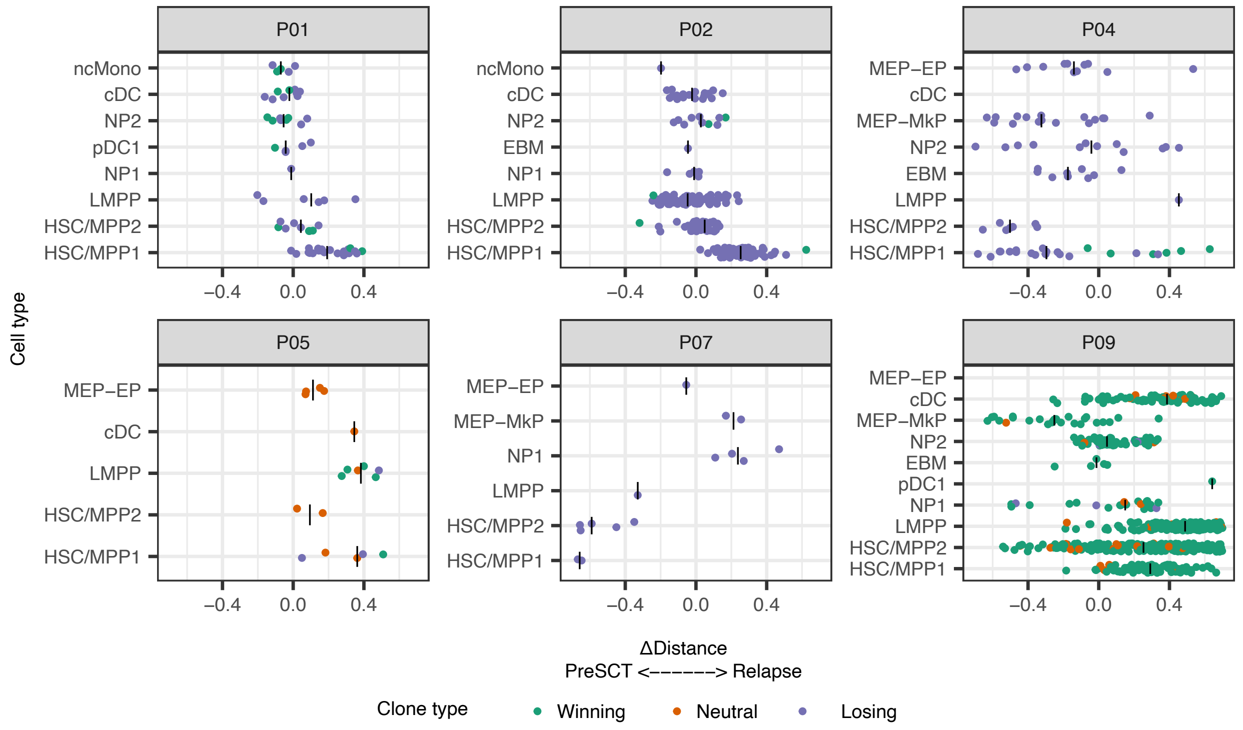


**Fig. S15 Deviation of individual CD34⁺ uMRD cells toward bulk preSCT or relapse states across hematopoietic progenitors.** Deviation scores of individual CD34⁺ uMRD cells across cell types in patients P01, P02, P04, P05, P07, and P09. Scores quantify the phenotypic similarity of each uMRD cell to preSCT versus relapse states. Positive values indicate greater similarity to relapse, while negative values indicate greater similarity to preSCT. Points represent single cells and are colored by clonal status (green, winning; orange, neutral; purple, losing).

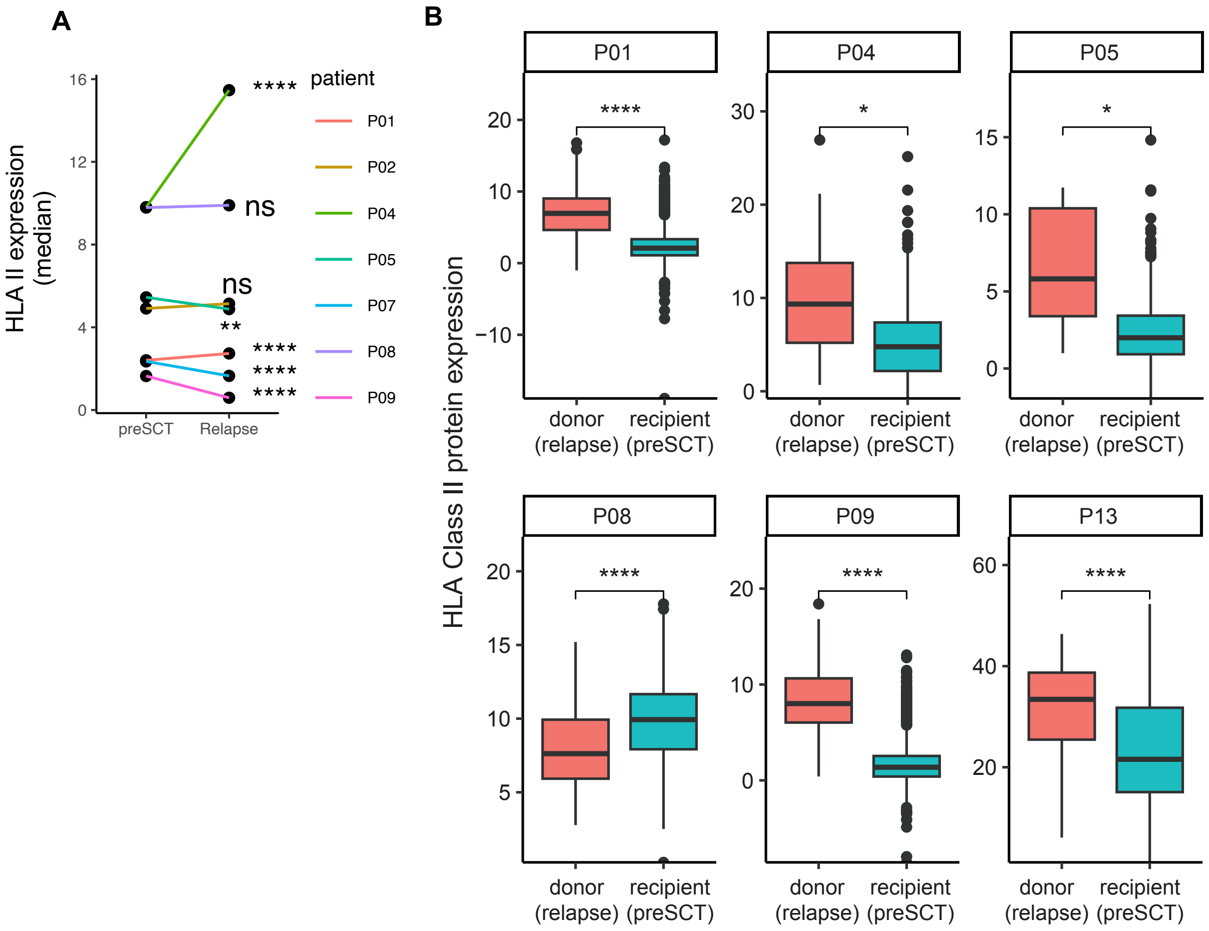


**Fig. S16**. **HLA class II protein expression in MDS.** (A) Boxplots show HLA class II protein expression (CITE-seq) in donor cells from relapse samples and recipient (malignant) cells from pre-SCT samples. To control for CITE-seq protein batch effects, comparisons were restricted to cells processed within the same batch. Center lines indicate medians; boxes, interquartile ranges; whiskers, 1.5× IQR; points, outliers. (B) Paired dot plot of median HLA class II expression in malignant cells from pre-SCT and relapse samples across patients, with lines connecting samples from the same patient. Statistical comparisons were performed using the Wilcoxon rank-sum test, and significance levels are indicated (*P < 0.05; **P < 0.01; ***P < 0.001; ****P < 0.0001; ns, not significant).

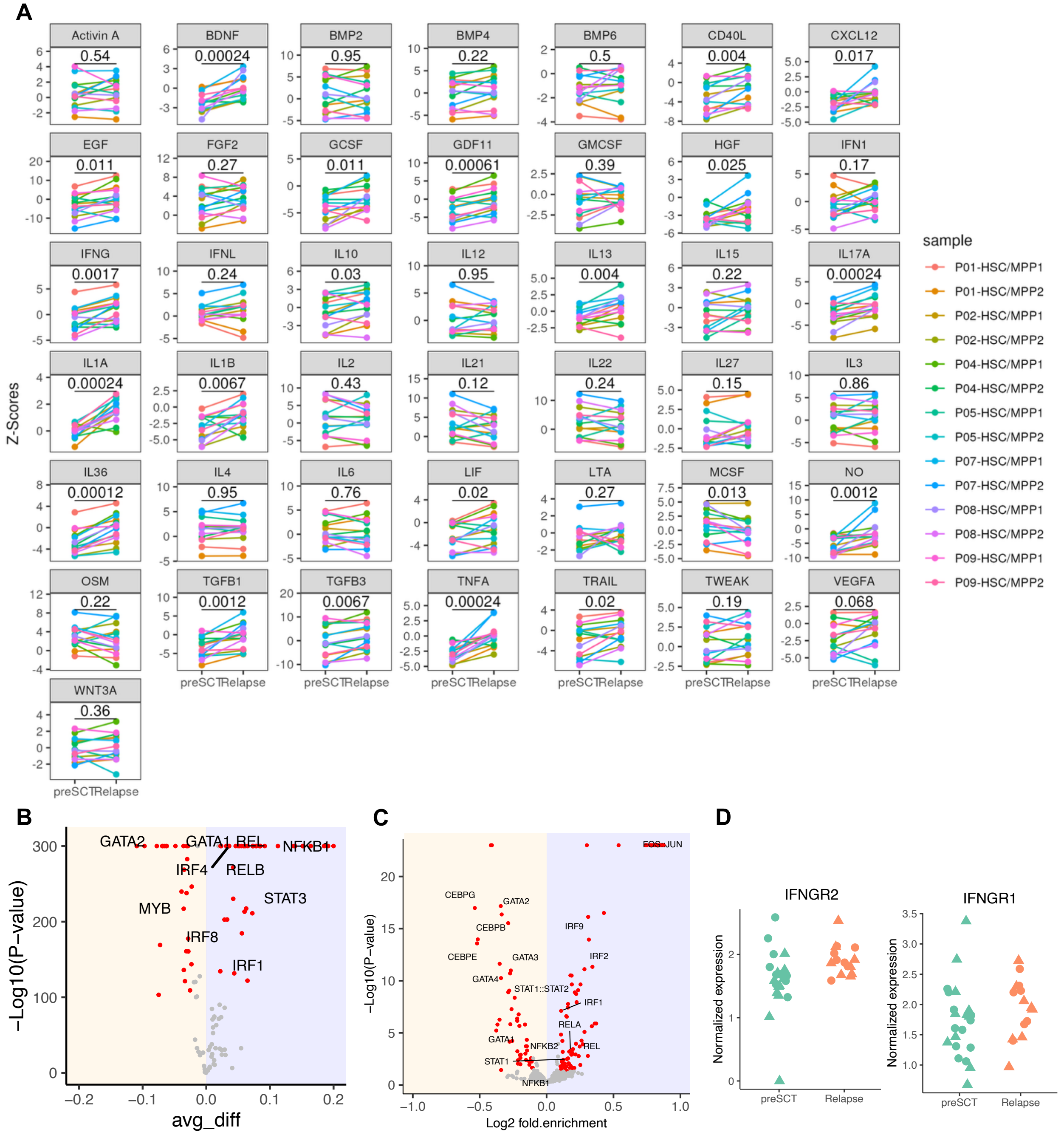

**Fig. S17. Altered cytokine signaling in HSC/MPP cells at relapse**. (A) Cytokine signaling activity (Z-scores) was inferred using CytoSig from pseudobulk gene expression profiles. Dynamics are shown for matched preSCT and relapse HSC/MPP subsets across patients, with lines connecting paired samples; P values were calculated by paired Wilcoxon tests. (B) Differential eRegulon activity between relapse and preSCT HSC/MPP cells across patients. (C) Motif enrichment analysis on differential peaks comparing relapse versus preSCT HSC/MPP cells. (D) Normalized expression of IFNGR2 and IFNGR1 in preSCT and relapse HSC/MPP cells.
